# A noncanonical polyamine from bacteria antagonizes animal mitochondrial function

**DOI:** 10.1101/2024.04.29.591726

**Authors:** Kelsie M. Nauta, Darrick R. Gates, Matthew Weiland, Marco E. Mechan-Llontop, Xiao Wang, Kim P. Nguyen, Christine Isaguirre, Megan R. Genjdar, Ryan D. Sheldon, Connie M. Krawczyk, Nicholas O. Burton

## Abstract

Canonical polyamines such as agmatine, putrescine, and spermidine are evolutionarily conserved metabolites found in nearly all forms of life ranging from bacteria to humans. Recently, interactions between polyamines produced by gut bacteria and human intestinal cells have been proposed to contribute to both Irritable Bowel Syndrome with Diarrhea (IBS-D) and inflammatory bowel diseases. However, the molecular mechanisms that underlie these effects are often unclear due in part to limitations in the methods used to manipulate and study polyamine functions *in vivo.* Here, we developed a *Caenorhabditis elegans* based screening platform and a modified LC-MS approach for profiling polyamine metabolites. We combined these methods to make the unexpected discovery that dysfunctional polyamine metabolism in both Gram-negative (*E. coli*) and Gram-positive (*B. subtilis*) bacteria can result in the accumulation of a noncanonical polyamine intermediate, N^1^-Aminopropylagmatine (N^1^-APA). We further find that N^1^-APA is produced via spermidine synthase (SpeE) and that it is bioactive when encountered by animals. Specifically, we find that when N^1^-APA is produced by bacteria in animal intestines it can be transported into intestinal cells via the polyamine transporter CATP-5 where it antagonizes both animal development and mitochondrial function across diverse animal species. Lastly, we find that N^1^-APA functions analogously to the deoxyhypusine synthase inhibitor GC7. For example, like GC7, N^1^-APA antagonizes eIF5A hypusination and inhibits the alternative activation of mammalian macrophages. To our knowledge, these findings are the first to demonstrate that N^1^-APA is a bioactive metabolite and that bacteria can produce a small molecule that functions similarly to existing deoxyhypusine synthase inhibitors. Furthermore, these results suggest an exciting new mechanistic hypothesis for why the loss of *speB* in gut microbes, including *E. coli,* has been both linked to inflammatory bowel disease (IBD) in humans and found to drive IBD in germ free mice.

## Introduction

Canonical polyamine metabolites including agmatine, putrescine, spermidine, and spermine are found in nearly all domains of life and are generally present at millimolar levels in animal cells ^1,2^. Changes in the metabolism or abundance of polyamines have been linked to numerous human pathologies. For example, mutations in the polyamine transporter ATP13A2 cause a heritable form of juvenile onset Parkinson’s disease ^3–5^. Similarly, changes in the production of polyamines by gut microbiota have recently been found to both cause irritable bowel syndrome (IBS-D) and, in other contexts, have been proposed to protect against the development of inflammatory bowel diseases and colitis ^6,7^. Despite their clear importance to biology and demonstrable relevance to numerous human pathologies, the molecular mechanisms by which altered polyamine metabolism causes changes in animal physiology are often unclear. For example, polyamines have been proposed to function as signaling molecules ^8–10^ or antioxidants ^11,12^, to stabilize RNA, protein, and membranes ^13,14^, and to regulate mRNA translation via the hypusination of the transcription factor, eIF5A^15^. However, in many cases the exact molecular mechanism by which changes in polyamine metabolism cause animal disease remain unknown. In addition, there are several cases of seemingly conflicting models for the mechanisms by which polyamines regulate animal physiology. For example, using the polyamine analog inhibitor of deoxyhypusine synthase, GC7, Puleston et al^16^ and Nakamura et al.^6^ have proposed that polyamines regulate alternative activation of macrophages (M2-like) through regulating the hypusination of eIF5A which in turn regulates the translation of a subset of mitochondrial proteins. By contrast, using a genetic approach, Anderson-Baucum et al.^17^ reported no clear evidence that hypusination of eIF5A regulated mitochondrial protein abundance by proteomics and suggested that previous findings might be due to alternate target effects of GC7. Instead, they found that eIF5A hypusination levels decreased the proinflammatory nature of macrophages by decreasing translation of proinflammatory mRNAs^17^, Thus, the precise mechanisms by which polyamines regulate aspects of animal physiology such as immune cell activation remain unclear.

Studies describing how the disruption of polyamine metabolism causes or contributes to disease in animals face multiple challenges. First, manipulating polyamine levels *in vivo* is challenging because the loss of enzymes required to synthesize putrescine and spermidine are lethal in many animals. In animal intestines, polyamines can also be transported between intestinal cells and gut microbes, complicating the interpretation of animal studies due to potential compensatory effects ^2,18^. Despite these challenges, recent studies have found that when agmatinase (encoded by *speB*) is deleted from bacterial genomes that inhabit animal intestines, mice develop colitis and *C. elegans* undergo developmental arrest ^6,7^. SpeB converts agmatine to putrescine in the canonically polyamine synthesis pathway^19^, and both excess agmatine and loss of putrescine have been proposed as the underlying molecular mechanism to explain these changes to animal physiology; however, neither has been shown to be sufficient to cause these effects ^6,7^.

The ability to measure polyamine levels *in vivo* also faces technical difficulties. For example, most kit-based approaches only quantify total polyamines ^20^. To overcome this, mass spectrometry-based approaches have been employed. However, polyamines are poorly detected in standard metabolomics-based approaches ^21^. Additionally, while mass spectrometry methods have been developed to reliably quantify the core polyamine metabolites (agmatine, putrescine, and spermidine), most studies are unable to simultaneously identify the many variants of these molecules. Thus, studies of polyamines are largely restricted to quantifying canonical polyamines and might overlook alternative polyamine metabolites. As a consequence of these limitations, defining cause-and-effect relationships between polyamines and mRNA, protein, or membrane stability, remains difficult. Thus, the precise molecular mechanisms for how disruptions in polyamine metabolism lead to human diseases remain unclear.

While screening a complete non-essential *Bacillus subtilis* gene knockout library for bacterial genes that regulate animal development in *C. elegans*; we discovered that the loss of *speB* in *B. subtilis* caused complete animal developmental arrest. Two previous studies found the loss of *speB* in *E. coli* results in the failure of *C. elegans* to develop in the presence of excess agmatine or causes mouse lethality in models of inflammatory bowel disease ^6,7^. Unexpectedly, however, we found that the effects of *speB* mutant *B. subtilis* on animals were not due to a requirement for bacterial putrescine in animals or due to excess agmatine. Instead, we found that diverse Gram-negative and Gram-positive bacteria produce a non-canonical polyamine (N^1^-Aminopropylagmatine (N^1^-APA)) via the activity of the conserved bacterial enzyme SpeE on agmatine, corroborating recent biochemical findings ^22,23^. In addition, we found that N^1^-APA accumulates in *speB* mutant bacteria and is transported into animal intestinal cells via the polyamine transporter CATP-5. Once inside animal cells, N^1^-APA acts analogously to GC7, a chemical inhibitor of deoxyhypusine synthase. These findings demonstrate that N^1^-APA functions like GC7 in both invertebrates and vertebrates which has numerous downstream consequences, such as inhibiting *C. elegans* development and immune cell activation in mouse bone marrow-derived macrophages (BMM_Φ_). To our knowledge, these findings are the first to report that N^1^-APA is a bioactive metabolite in any species and the first observation that a polyamine metabolite derived from bacteria functions analogously to the deoxyhypusine synthase inhibitor GC7 *in vivo*. These findings demonstrate the importance of investigating individual non-canonical polyamines when studying how changes in polyamine metabolism cause disease. They also provide an alternative N^1^-APA dependent mechanism of action that may explain why recent reports link the loss of *speB* in gut microbiome bacteria to inflammatory bowel disease in both mice and humans ^6,24^.

## Results

### Loss of *speB* in bacteria antagonizes animal development

To identify new links between bacterial metabolism and animal physiology, we screened a complete *B. subtilis* non-essential gene knockout library for changes in animal development, using *Caenorhabditis elegans* as a model (Fig. 1A – See methods for screen details) ^25^. From our screen of 3,984 mutant *B. subtilis* isolates, we identified only a single *B. subtilis* gene, *speB*, that was required for *C. elegans* to develop to adulthood (Fig 1B). SpeB encodes an agmatinase which plays an intermediate role in the synthesis of spermidine (Fig. 1C) ^19,26^. We confirmed that *speB* is required for animals to develop to adulthood by complementing this mutant with a wild-type copy of *speB* (Fig. 1B). We conclude that *speB* is required in *B. subtilis* to promote *C. elegans* development to adulthood.

**Figure 1:**
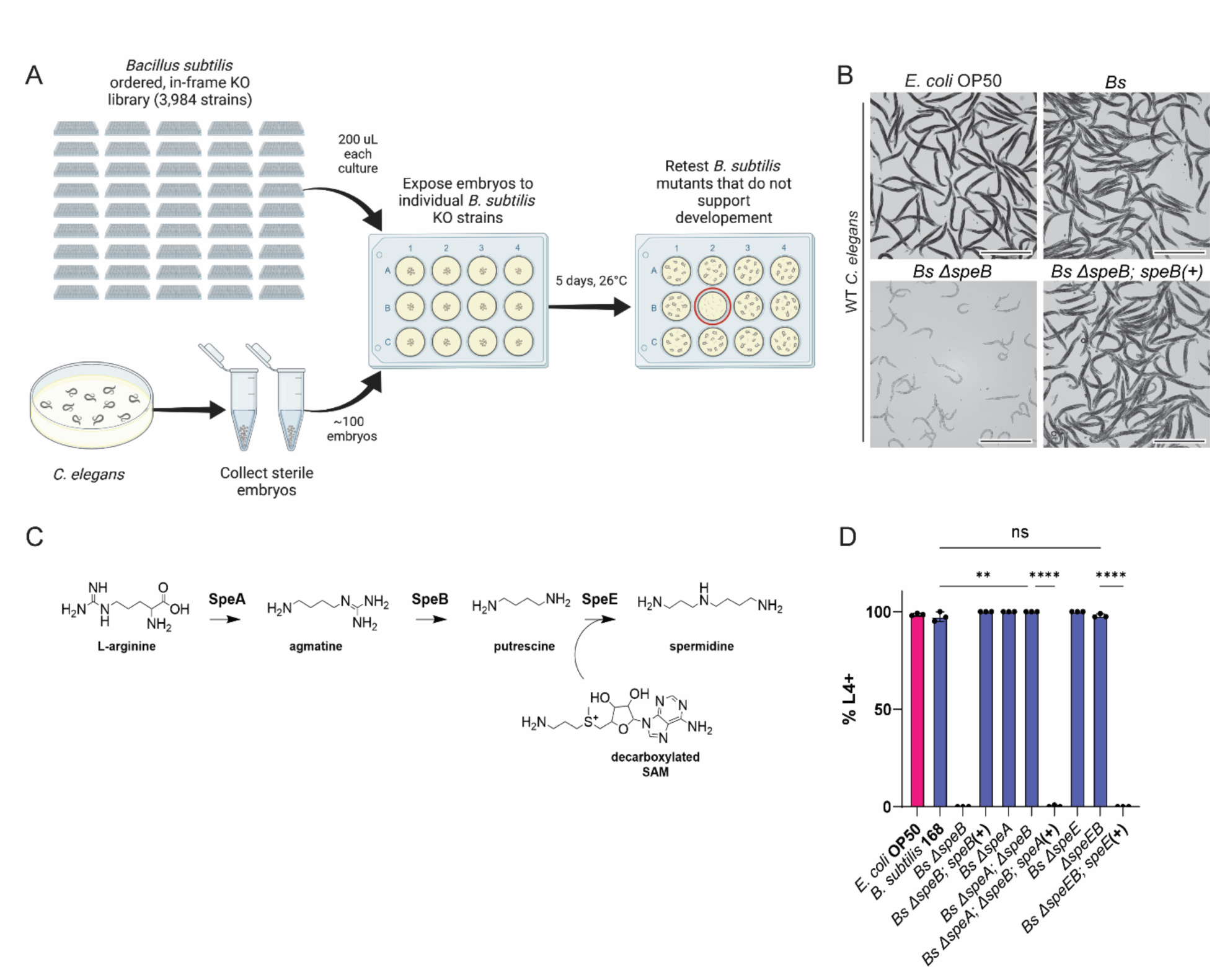
Loss of *speB* in bacteria antagonizes animal development. (A) Graphical representation of the platform used to screen for mutants of *B. subtilis* that did not support animal development. Germ-free embryos were arrayed on the library of 3,984 *B. subtilis* mutants and screened for development beyond the L4 stage. Candidate mutants were retested on 60 mm plates with ∼500 embryos. (B) Representative images of WT *C. elegans* on *E. coli* OP50, *B. subtilis* 168 (*Bs*)(NOBb513), *Bs* Δ*speB* (NOBb514), and *Bs* Δ*speB; speB*(+) (NOBb475). The animals were incubated at 20 °C for 3 days on their respective diets, washed in M9 buffer, and paralyzed with tetramisole (∼10 µg/mL). The scale bar is 1 mm. (C) Canonically, in *B. subtilis*, spermidine is synthesized from L-Arginine by the consecutive activities of SpeA, SpeB, and SpeE. (D) Percentage of WT *C. elegans* that developed pasts L4+ on diets of *Bs* Δ*speE* (NOBb517), *Bs* Δ*speEB* (NOBb515), *Bs* Δ*speEB; speE* (+)(NOBb501), *Bs* Δ*speA* (NOBb516), *Bs* Δ*speB*; Δ*speA* (NOBb518), and *Bs* Δ*speA*; Δ*speB*; *speA*(+) (NOBb503).The animals were incubated at 26 °C for 5 days on their respective diets and L4+ animals were quantified. Experiments were performed in triplicate with ∼500 animals per replicate. Error bars = s.d., **** = p < 0.0001, ** = p < 0.0065, ns = not significant

In many bacteria, SpeB (agmatinase) is part of a pathway, together with SpeA and SpeE (spermidine synthase), that converts arginine to spermidine^19^ (Fig. 1C). Previous studies have proposed that deletion of *speB* causes either the loss of beneficial downstream polyamines (putrescine, spermidine) or a toxic buildup of agmatine and that these changes in canonical polyamines results in deleterious effects on animal physiology^6,7^. To determine if *C. elegans* are unable to develop to adulthood on *B. subtilis* Δ*speB* because they require spermidine from their diet, we tested if the deletion of *speA* or *speE* in *B. subtilis* also caused animals to arrest their development (SFig. 1A). In contrast, we found that deletion of *speA* and *speE* did not alter WT animal development (Fig. 1D, SFig. 1B). Since *speA* converts L-arginine into agmatine^19^, we generated a double Δ*speA* Δ*speB* deletion strain to test if preventing the aberrant accumulation of agmatine prevented animal developmental arrest. We found deletion of *speA* in the Δ*speB* background rescued *C. elegans* development, suggesting agmatine causes development arrest (Fig. 1D, SFig. 1B). We confirmed this phenotype by complementing Δ*speA* Δ*speB* with *speA* (Fig. 1D, SFig. 1B). Surprisingly, we found that knocking out the *speEB* operon does not phenocopy Δ*speB* and that Δ*speEB* mutants behave similarly to Δ*speA* Δ*speB* mutant bacteria (Fig. 1D, SFig. 1B). We confirmed this result by complementing Δ*speEB* with *speE* and found that the strain no longer promoted animal development (Fig. 1D, SFig. 1B). We conclude that the deletion of *speB*, but not other genes in the spermidine synthesis pathway, antagonizes animal development and that this effect is dependent on both *speA* and *speE* activity. These findings suggest that the simple loss of spermidine or a buildup of agmatine are not the cause of Δ*speB-mediated* disruption of animal development.

### Loss of *speB* causes diverse species of bacteria to accumulate N^1^-APA

We hypothesized that *B. subtilis speB* mutants might produce a noncanonical polyamine that antagonizes animal development. This would be in line with recent reports that bacteria produce numerous noncanonical polyamines, sometimes using the same enzymes predicted to be involved in canonical polyamine metabolism, such as SpeE ^22^. To test this, we developed an untargeted LC-MS method for the profiling of polyamine metabolites in *B. subtilis* Δ*speA*, Δ*speB*, Δ*speEB*, Δ*speE*, Δ*speB*; Δ*speA*, and WT cultures. Polyamines are not well retained or ionized in typical LC-MS metabolomics approaches ^21^. So, to do this we generated carbamyl derivatives of metabolite extracts and performed untargeted metabolomics. Using this approach, we detected 2641 blank- and peak quality-filtered LC-MS features. Differential abundance analysis revealed a consistent feature, 388.29207 m/z at retention time of 3.781min, that was the most significantly increased compound in Δ*speB* cells versus the other polyamine synthesis mutants (Fig 2A). Evaluation of extracted ion chromatograms (EIC) indicated that this compound was only detectable in the Δ*speB* cells (Fig 2B). Using accurate mass MS1 measurements, we found three possible chemical formulae within 5ppm (Fig 2C). Of these, only one contained oxygen atoms, of which there are two per carbamyl derivative (C5H9O2; Figure 2C). From this we deduced that the unknown compound contained two carbamyl-derivatized sites, and subtraction of 2x(C5H9O2) from the predicted chemical formula of the unknown (C18H37N5O4) yielded a parent compound formula of C8H21N5. This is consistent with the non-canonical polyamine, N^1^-APA, that was originally reported to accumulate in *speB* mutant *Thermus thermophilus* ^22,23^ (N^1^-APA; (Figure 2D). To confirm this putative identification, we spiked in a synthetic standard to Δ*speB* media and observed an increase in the 388.2921 EIC over unspiked Δ*speB* controls (Figure 2E). MS2 fragmentation spectra of the endogenous unknown and the synthetic standard further confirmed the identity of the Δ*speB* specific polyamine as N^1^-APA (Figure 2F).

**Figure 2.**
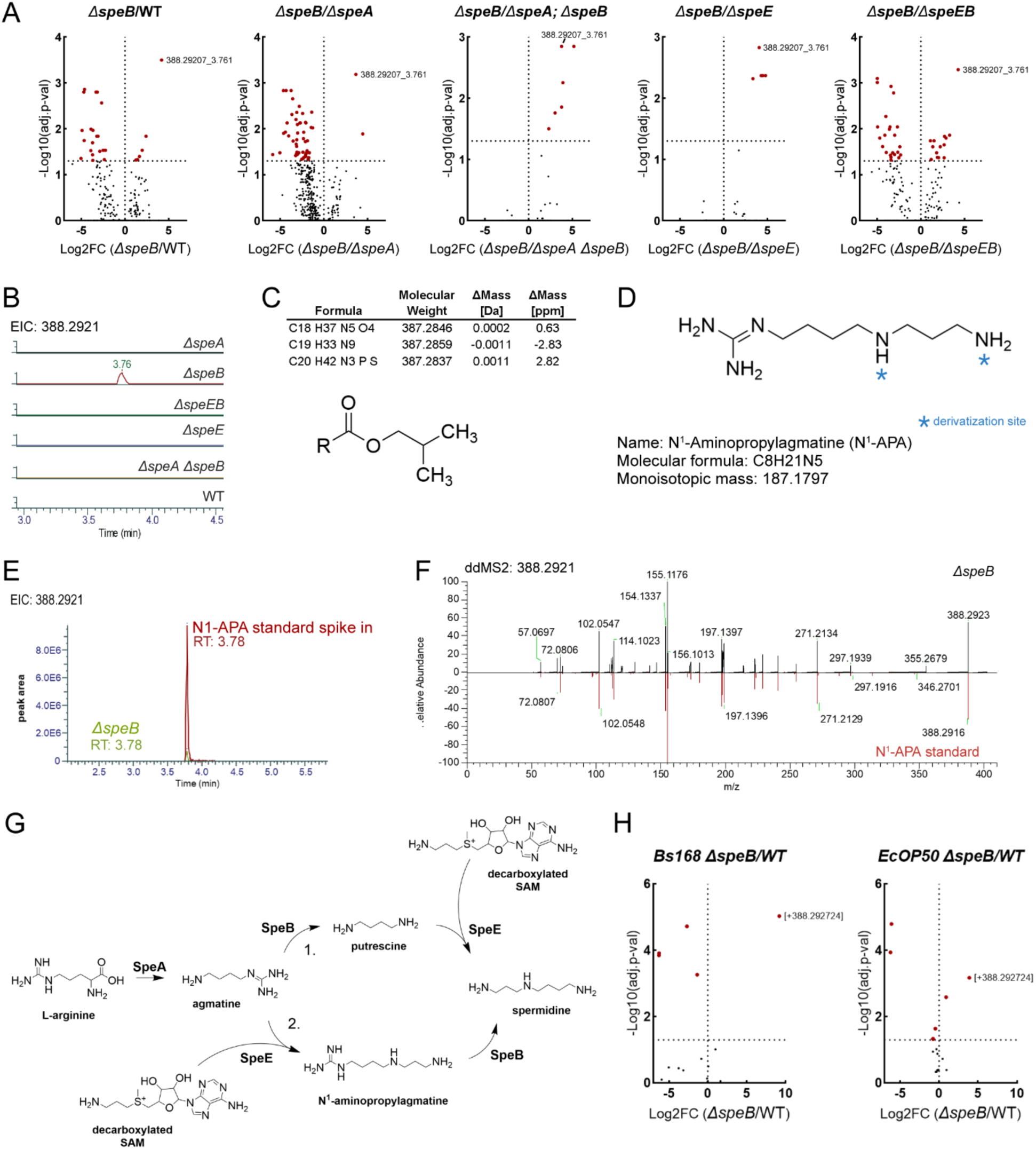
Identification and validation of N^1^-APA accumulated in Δ*speB* mutant bacteria. (A) Volcano plots showing differential abundance analysis of untargeted metabolomics on carbamylated-derivatives between Δ*speB* cell lysates and the other strains evaluated. Red dots are LC-MS features with an FDR adjusted P-value <0.05. Note the feature 388.29207 m/z at retention time 3.781min is the most significantly elevated compound in each comparison. (B) Representative extracted ion chromatograms of 388.29207 m/z (+/- 5ppm). (C) Top, chemical formula predictions for m/z 388.29207 using accurate mass and fine isotopic structure in Compound Discoverer. Bottom, structure of carbamylated-derivatives indicate two oxygens are present in each derivatized moiety. (D) Structure of N^1^-APA with sites of carbamylated-derivatives denoted with a blue asterisk. Note the chemical formula of N^1^-APA (C8H21N5) with carbamylated-derivatives (C5H9O2 x2) at the indicated sites match the predicted chemical formula of the unknown (C18H37N5O4). (E) Overlaid extracted ion chromatograms of Δ*speB* and Δ*speB*+N1APA spike-in provides accurate mass and chromatographic profile confirmation of N1APA as the unknown feature 388.29207@3.7981min. (F) Data dependent MS2 fragmentation mirror plot of parent ion 388.29207@3.7981min in Δ*speB* (top) and neat N^1^-APA standard (bottom) provides further confirmation that the unknown is N^1^-APA. (G) Canonical spermidine synthesis pathway (1.) compared to the alternative pathway (2.) found in *Thermus thermophilus*, which includes N^1^-APA as an intermediate metabolite. (H) Volcano plots showing differential abundance analysis of targeted polyamine carbamylated-derivatives in cell lysates between Δ*speB* and other evaluated lines. The feature identified as N^1^-APA is elevated in *E. coli* OP50 Δ*speB* as compared to WT *E. coli* OP50. Both strains were grown in the presence of 50 mM agmatine.

Previous studies of *C. elegans* found that the loss of *speB* in *E. coli* also caused animals to arrest their development, similar to our findings for *B. subtilis*^7^. Similar to studies in *C. elegans*, studies in germ free mice found that *speB* mutant *E. coli* (but not wild-type *E. coli*) alter mouse physiology and drive inflammatory bowel disease-like pathologies in mammals. Collectively, these studies suggest that the loss of *speB* in other species of bacteria also affects animal physiology in taxa ranging from nematodes to mammals. Despite these consistent findings for *speB* mutant *E. coli* affecting many important aspects of animal physiology, the mechanistic explanation remains unclear with different studies proposing the effects might be due to a buildup of agmatine or a loss of putrescine. Our findings from *B. subtilis* suggest that in addition to any potential effects on canonical polyamines (agmatine, putrescine), *speB* mutant *E. coli* might also accumulate the non-canonical polyamine N^1^-APA because it disrupts a proposed alternative spermidine synthesis pathway (Fig. 2G). To test this, we profiled the abundance of polyamines, including N^1^-APA, in *E. coli* with and without *speB*. As in *B. subtilis*, we found that *E. coli* lacking *speB* also accumulated N^1^-APA (Fig. 2H). We conclude that *speB* mutant *E. coli* also accumulate N^1^-APA. Given that Gram positive (*B. subtilis*) and Gram negative (*E. coli*) bacteria diverged >2 billion years ago, our findings suggest that most bacteria may produce N^1^- APA in certain contexts. We note that this hypothesis would also be consistent with a recent biochemical study that found that 17 out of the 18 tested bacterial spermidine synthases expressed in *E. coli* BL21 (including *speE* from *B. subtilis*) were able to convert agmatine into N^1^-APA, similar to the original report of N^1^-APA production in *T. thermophilus* ^22,23^.

### N^1^-APA is a bioactive metabolite that drives mitochondrial dysfunction and developmental arrest in animals when transported via the polyamine transporter CATP-5

Our previous genetics studies of *B. subtilis* indicated that the effects of *speB* mutant *B. subtilis* on *C. elegans* development were not due to either the loss of putrescine or the accumulation of agmatine (Fig. 1). Given our mass spectrometry findings, we hypothesized that the accumulation of N^1^-APA in *speB* mutant bacteria might explain why *C. elegans* arrest their development in response to *speB* mutant *B. subtilis* (Fig. 2). To test this, we synthesized N^1^-APA and grew *C. elegans* on a diet of WT *B. subtilis* on media containing exogenous N^1^-APA. We found exogenous N^1^-APA was sufficient to cause developmental arrest in *C. elegans* (Fig. 3A). This developmental arrest was not observed when spermidine or agmatine were supplemented into plates (Fig. 3A, SFig. 2). We also found the addition of exogenous spermidine to the media is sufficient to suppress the effects of exogenous N^1^-APA (Fig. 3B) or *B. subtilis ΔspeB* (Fig. 3C). We conclude that N^1^-APA production by *B. subtilis* antagonizes animal development, possibly by disrupting the normal function of spermidine.

**Figure 3:**
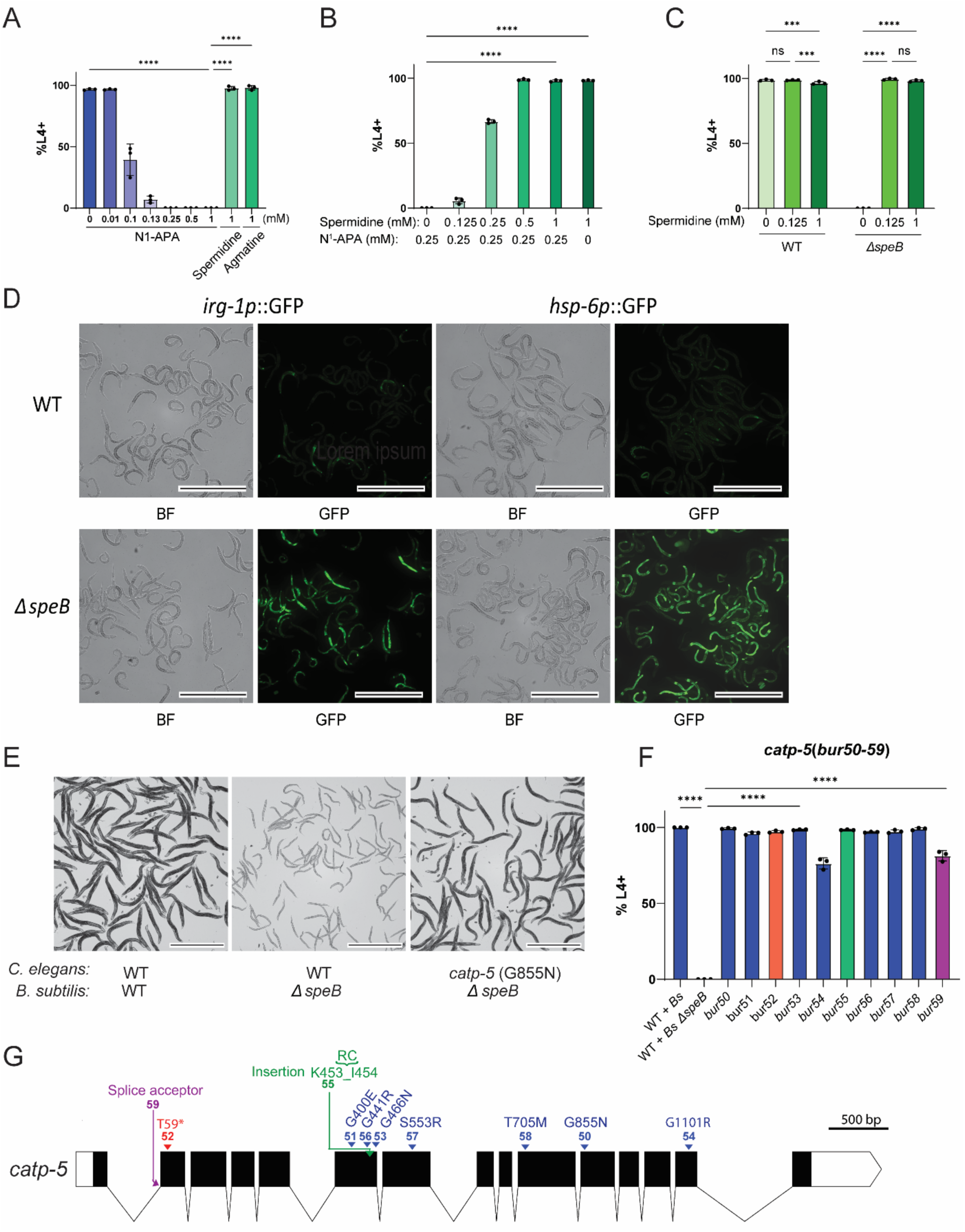
Mutations in the polyamine transporter, *catp-5*, confer resistance to *B. subtilis* Δ*speB*. (A) Percent of WT *C. elegans* (N2) that developed past L4 when grown on media supplemented with a range of concentrations of N^1^-APA and seeded with wild type *B. subtilis* after incubating for 3 days at 20 °C. Control plates were supplemented with 1 mM agmatine and spermidine. Experiments were performed in triplicate with ∼200 animals per replicate. Error bars = s.d. **** = p < 0.0001. (B) Percent of WT *C. elegans* that develop past L4 on modified NGM (see methods) supplemented with 0.25 mM N^1^-APA and a concentration gradient of spermidine (0.125 —1 mM). Experiments were performed in triplicate with ∼300 animals per replicate. Error bars = s.d. **** = p < 0.0001. (C) Percent of WT *C. elegans* that developed past L4 on modified NGM (see methods) seeded with *B. subtilis* (WT) or *B. subtilis ΔspeB* (NOBb369) and supplemented with spermidine (0.125 and 1 mM). Experiments were performed in triplicate with ∼300 animals per replicate. Error bars = s.d., *** = p < 0.008, **** = p < 0.0001, ns = not significant. (D) Brightfield (BF) and fluorescent (GFP) images of growth stage matched *irg-1p::GFP* and *hsp-6p::GFP* on a diet of *B. subtilis* (NOBb513) or *B. subtilis ΔspeB* (NOBb514). The *C. elegans* strains were incubated on the respective *B. subtilis* strains for ∼30 hours at 20 °C. Representative images were pseudo-colored using ImageJ v1.54f. Scale bars are 0.5 mm. (E) Representative images of WT *C. elegans* and mutant *catp-5*(*bur50*) on *B. subtilis* 168 (*Bs*)(NOBb513) and *B. subtilis* 168 Δ*speB* (NOBb514). The animals were incubated at 20 °C for 3 days on their respective diets, washed in M9 buffer, and paralyzed with tetramisole. The scale bar is 1 mm. (F) Percent of WT *C. elegans* (N2) and *catp-5*(*bur50-59*) that developed past L4 on *B. subtilis* 168 Δ*speB* (NOBb514) after 5 days at 26 °C. WT *C. elegans* on *B. subtilis* 168(*Bs*)(NOBb513) was included as a control. Experiments were performed in triplicate with ∼500 animals per replicate. Error bars = s.d., **** = p < 0.0001 (G) Graphical representation of mutations in CATP-5 that restored animal development per allele (*bur50-59*). Red = nonsense mutation. Blue = nonsynonymous mutation. Green = insertion mutation. Purple = splice adaptor mutation. Numbers 50-59 correspond to allele *bur50-bur59*.

To determine how N^1^-APA antagonizes animal development, we screened 11 *C. elegans* GFP stress reporters on a diet of either *B. subtilis* WT or *B. subtilis ΔspeB* (Table 2). We found that GFP reporters for *hsp-6* and *irg-1* were both substantially increased when animals were fed Δ*speB* mutant *B. subtilis* when compared to WT *B. subtilis* (Fig. 3D) or compared to other polyamine synthesis mutants (SFig. 3). *hsp-6* expression is part of the mitochondrial unfolded protein response and is activated in response to mitochondrial stress. *irg-1* was originally used as a reporter for bacterial infections, but was later found to be activated in response to mitochondrial stress^27^. Thus, the activation of both *hsp-6* and *irg-1* is consistent with animals experiencing mitochondrial stress. This activation is reversed when a wild-type copy of *speB* was restored to the *B. subtilis* genome (SFig. 3). Importantly, we did not observe a difference in level of fluorescence for the other nine reporters that were tested: *nlp-29, clec-60, gcs-1, hsp-60, hsp-16.2, gst-4, hsp-4, sod-3, bvIs5* (Table 2). The activation of the mitochondrial unfolded protein response is likely not due to general proteostasis stress because we did not observe activation of *hsp*-*4::*GFP which is activated as part of the endoplasmic reticulum unfolded protein response (SFig.4, Table 2). We conclude that N^1^-APA activates the *C. elegans* mitochondrial unfolded protein response, likely by causing mitochondrial stress.

**Figure 4:**
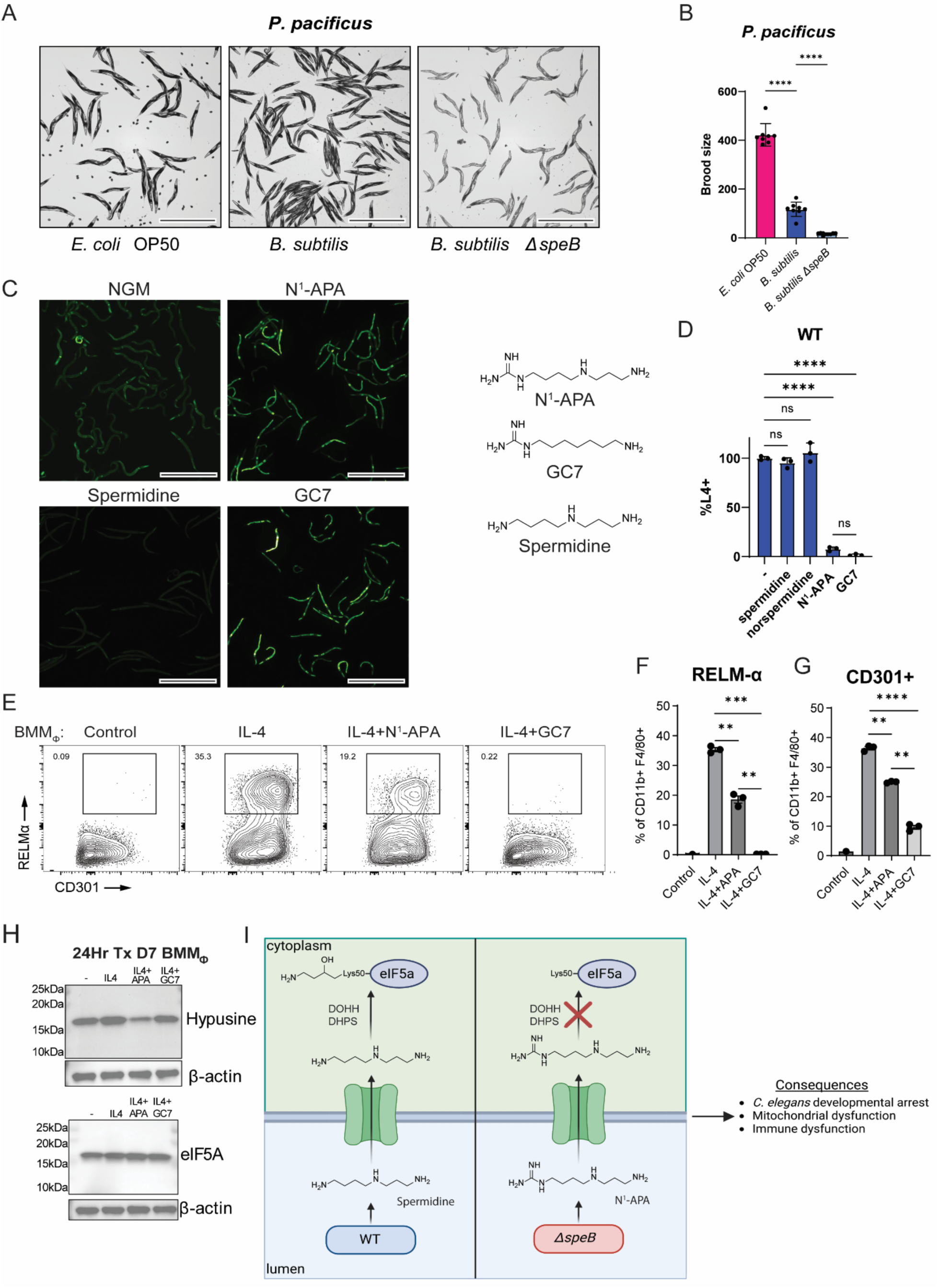
N^1^-APA functions similarly to the deoxyhypusine synthase inhibitor GC7 in diverse organisms. (A) Representative images of *Pristionchus pacificus* on a diet of *E. coli* OP50, *B. subtilis*, and *B. subtilis* Δ*speB* (NOBb369). Scale bars are 1 mm. (B) Quantification of brood size in *P. pacificus* on a diet of *E. coli* OP50, *B. subtilis*, and *B. subtilis* Δ*speB* (NOBb369). Error bars = s.d., **** = p < 0.0001. (C) Fluorescent (GFP) images of growth stage matched *C. elegans* encoding *hsp-6p::GFP* fed a diet of *B. subtilis* (NOBb513) on NGM agar plates supplemented with 0.125 mM spermidine, N^1^-APA, and GC7. Chemical structures are shown to the right. The *C. elegans* strain were incubated with *B. subtilis* strain for ∼30 hours at 20 °C. Representative images were pseudo-colored using ImageJ v1.54f. Scale bars are 0.5 mm. (D) Percent of WT *C. elegans* that developed past L4 on *B. subtilis* 168 (NOBb513) after exposure to 0.125 mM spermidine, norspermidine, N^1^-APA, and GC7 for 3 days at 20 °C. Experiments were performed in triplicate with ∼500 animals per replicate. Error bars = s.d., **** = p < 0.0001. ns = not significant. (E) Representative flow plots, gates of RELMα+ BMM_Φ_. Proportions of (F) RELMα and (G) CD301 BMM_Φ_ (IL-4) following treatment with 200 µM N^1^-APA or GC7 for 24 hours. Experiment performed in triplicate; representative of 2 independent experiments. Error bars = S.E.M., **** = p < 0.0001, *** = p < 0.009, ** = p < 0.01. (H) N^1^-APA inhibits eIF5a hypusination in BMM_Φ_. Representative images of 3 independent experiments. (I) N^1^-Animopropylagmatine accumulates in the *C. elegans* intestinal lumen when the animals are grown on a diet of *B. subtilis ΔspeB*. CATP-5 transports N^1^-APA into intestinal cells where it inhibits eIF5a hypusination. N^1^-APA antagonizes mitochondrial and immune function.

To determine how a diet of *B. subtilis ΔspeB* antagonizes animal development, we mutagenized wild-type *C. elegans* and screened for mutant animals that could develop to adulthood when feeding on *B. subtilis ΔspeB.* We isolated 10 lines of mutant animals that developed to adulthood even when fed *B. subtilis ΔspeB* (Fig. 3E, 3F, SFig. 5, Table 1). We found the mutant lines harbored predicted loss-of-function mutations in the *catp-5* polyamine transporter (Fig. 3G). CATP-5 is expressed in intestinal cells and was previously reported to import polyamines from the intestinal lumen into the intestinal cells ^28^. These findings are consistent with a model in which N^1^-APA is produced by *B. subtilis ΔspeB* and then is imported by CATP-5 into animal intestinal cells where it antagonizes animal development.

**Table 1:** List of mutations in *catp-5* that result in *C. elegans* development on *B. subtilis ΔspeB*.

|  | Position | Reference | Alternate | Variant Type | CDS Variant Annotation | Protein Variant Annotation | Alternate Allele Depth |
| --- | --- | --- | --- | --- | --- | --- | --- |
| *bur50* | 8092275 | C | T | missense_variant | c.4267G>A | p.Gly855Asp | 103 |
| *bur51* | 8094229 | C | T | missense_variant | c.2313G>A | p.Gly400Glu | 95 |
| *bur52* | 8095771 | C | T | stop_gained | c.770G>A | p.Trp59* | 106 |
| *bur53* | 8094031 | C | T | missense_variant | c.2510G>A | p.Gly466Asp | 112 |
| *bur54* | 8091401 | C | T | missense_variant | c.5141G>A | p.Gly1101Arg | 65 |
| *bur55* | 8095831 | C | T | splice_acceptor_variant | c.118-1G>A | intron | 78 |
| *bur56* | 8094107 | C | T | missense_variant | c.2435G>A | p.Gly441Arg | 103 |
| *bur57* | 8093719 | A | T | missense_variant | c.2822T>A | p.Ser553Arg | 93 |
| *bur58* | 8092772 | G | A | missense_variant | c.3770C>T | p.Thr705Met | 98 |
| *bur59* | 8094738 | G | A | insertion | c.1059+11C>T | p.Lys453-Iso454 in-sArgCys | 73 |

**Table 2:**
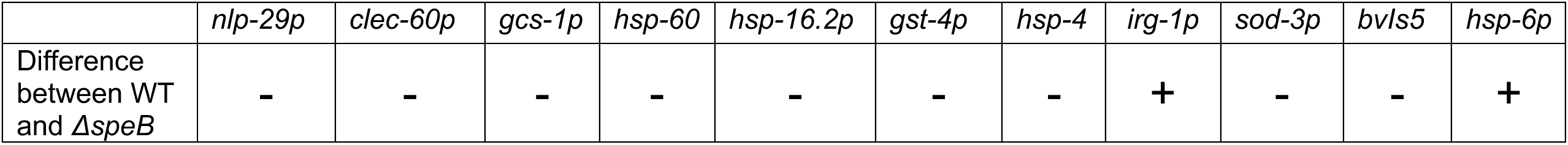
Summary of *C. elegans* GFP reporter strains on a diet of *B. subtilis 168 ΔspeB*.

### N^1^-APA functions analogously to the deoxyhypusine synthase inhibitor GC7 in both invertebrates and vertebrates

The effects of N^1^-APA could be specific to *C. elegans* or could generally antagonize animal development and mitochondrial function across species. To test this, we exposed the nematode *Pristionchus pacificus* to wild-type and *B. subtilis ΔspeB* (Fig.4A)*. P. pacificus* and *C. elegans* shared a last common ancestor approximately 100 million years ago ^29,30^. We found that *B. subtilis ΔspeB* also antagonized the development of *P. pacificus* and led to a >90% reduction animal fertility (Fig. 4B). We conclude that N^1^-Aminiopropylagmatine broadly affects nematode development and physiology and that this effect is not specific to *C. elegans*.

In the polyamine-eIF5A-hypusine axis, which is conserved in most animals from nematodes to humans, spermidine is used to hypusinate eIF5A and alter the translation of a subset of mRNAs. One of the key enzymes in this pathway is deoxyhypusine synthase which is also conserved from *C. elegans* to humans ^31^. Previous chemical inhibition studies of deoxyhypusine synthase found that GC7, a synthetic polyamine, was a potent inhibitor of deoxyhypusine synthase^16,32^. Furthermore, this study tested many derivatives of GC7 and found that N^1^-APA could also inhibit deoxyhypusine synthase, but that it was less potent than GC7 ^32^. At the time of this work, it was not known that bacteria synthesize N^1^-APA. Given that deoxyhypusine synthase is evolutionarily conserved, we hypothesized that N^1^-APA might cause both mitochondrial stress and developmental arrest in *C. elegans* by functioning analogously to GC7. To test this, we first exposed wild-type animals expressing the mitochondrial stress reporter *hsp-6::GFP* to either N^1^-APA or GC7 and found that *hsp-6::GFP* expression is activated by both molecules (Fig. 4C). We also found that both GC7 and N^1^-APA caused animals to arrest their development (Fig. 4D). Importantly, neither spermidine nor norspermidine activated *hsp-6::GFP* expression or caused developmental arrest at the relevant concentrations, suggesting these results are not broadly caused by excess exogenous polyamines (Fig. 4C, 4D, SFig. 6). We conclude that GC7 and N^1^-APA cause similar activation of a mitochondrial stress reporter and developmental arrest in *C. elegans*.

In mammals, polyamines are important for the regulation of macrophage activation. Activation of macrophages by IL-4 leads to an anti-inflammatory phenotype of macrophages (M2-like) that can be inhibited by GC7 and promoted by putrescine ^6,16,17^. Therefore, we tested whether N^1^-APA also altered the activation of mammalian macrophages in response to IL-4. Consistent with N^1^-APA being analogous to GC7 and previous studies^16^, we found N^1^-APA inhibits IL-4 induced expression of M2-like macrophage markers RELM-α and CD301 (Fig. 4E, 4F, 4G). While GC7 was more effective at inhibiting M2 phenotype than N^1^-APA, N^1^-APA treatment decreased hypusination of eIF5A to a greater extent than GC7 (Fig. 4H). These findings demonstrate that N^1^-APA inhibits the hypusination of eIF5A *in vivo* are consistent with a model in which N^1^-APA interferes with a normal function of spermidine in the polyamine-eIF5A-hypusine axis^16^.

## Discussion

In summary, our findings establish that N^1^-APA, produced via an alternative spermidine synthesis pathway in bacteria, can influence animal development and eIF5a hypusination when it is taken up by CATP-5 in animal intestinal cells (Fig. 4I). To our knowledge, these findings are the first to demonstrate that the bacterial polyamine intermediate, N^1^-APA, is bioactive in animals or that bacteria produce an inhibitor that functions similarly to the deoxyhypusine synthase inhibitor, GC7. Thus, we establish a novel link between a bacterial intermediate metabolite and the activity of a specific molecular pathway in animals. Our findings also further establish *C. elegans* as an excellent model species for identifying links between specific bacterial metabolites and target enzymes in animals.

Importantly, our findings suggest that the role of N^1^-APA in biology might be far more profound that previously appreciated. Specifically, our findings indicate that N^1^-APA not only is produced by both Gram-positive and Gram-negative species of bacteria, but that it can be produced at significant enough quantitates to alter animal development. Recently, Li *et al.,* 2024 reported the surprising finding that 17 out of 18 bacterial spermidine synthases, including *speE* from *B. subtilis*, can produce N^1^-APA when expressed in *E. coli*. These unexpected findings led the authors to conclude that “the *N*^1^-Aminopropylagmatine pathway for spermidine biosynthesis, which bypasses putrescine, may be far more widespread [in bacteria] than realized and may be the default pathway for spermidine biosynthesis in species encoding L-Arginine decarboxylase for agmatine production”. Why diverse species of bacteria might use this alternative non-canonical polyamine synthesis pathway (Fig. 2G) is unknown. However, our findings reported here indicate that N^1^-APA is a bioactive metabolite, that it inhibits nematode development, and that inhibits the activation of mammalian macrophages. Given these findings, we speculate that N^1^-APA might also function as an immune modulator in nature and/or function to modify animal physiology in the contentious ecosystem of soil microbiology. For example, non-canonical polyamine synthesis may act to protect soil microbes from being consumed by predatory animals such as nematodes. Future studies exploring how bacterial polyamines affect animal physiology and immune cell activation will further elucidate this relationship.

Our findings also suggest a potentially new mechanistic model for why previous studies found that disrupting *speB* in intestinal bacteria affects diverse aspect of animal physiology across both *C. elegans* and mammals. For example, a recent study of the effects of metformin on animal lifespans found that bacteria-derived agmatine underlies the effects of metformin on host metabolism and longevity in animals including *C. elegans*^7^. This was in part based on the finding that supplementing agmatine to *C. elegans* feeding on *E. coli* Δ*speB* resulted in a similar developmental arrest as we observed for *C. elegans* feeding on *B. subtilis* Δ*speB*. Because *speB* converts agmatine to putrescine in the canonical polyamine synthesis pathway, excess agmatine was hypothesized to accumulate in *speB* mutants and alter animal physiology. Here, we found there was not a significant difference in agmatine relative abundance between WT and *E. coli ΔspeB*. However, *E. coli* Δ*speB* did accumulate N^1^-APA in the presence of exogenous agmatine. Thus, our findings suggest a potentially alternative mechanistic model for why the loss of *speB* alters the effects of metformin on animals^7^; the loss of *speB* in *E. coli* leads to a buildup of N^1^-APA which alters mitochondrial function in animals. In this model, the effects of metformin, a CI inhibitor, may be altered in animals exposed to *speB* mutant bacteria because N^1^-APA accumulates in these bacteria and disrupts animal mitochondrial function^7^.

Similar to how loss of bacterial *speB* affects *C. elegans* physiology, studies in mice and humans have linked loss of *speB* in the gut microbiome to inflammatory bowel disease in mammals ^24^. For example, it was recently found that colonizing germ-free mice with *speB* mutant *E. coli* (as well as deleting a parallel polyamine pathway) promoted the development of severe colitis in mice^6^. These effects on inflammatory bowel disease and colitis were proposed to be due to a lack of putrescine. However, our data suggest N^1^-APA might also contribute to disease and could explain why some of the phenotypes caused by *speB* mutant *E. coli* in mice are similar to those caused by deoxyhypusine synthase inhibition by GC7. Specifically, our findings suggest that the reason both *speB* mutant *E. coli* and GC7 were found to drive colitis-like changes in mice or mouse cells might be because *speB* mutant *E. coli* produce N^1^-APA which functions analogously to GC7 and inhibits the development of anti-inflammatory M2-like macrophages (i.e. “M2-like” macrophages, characterized by the production of RELMa and CD301 (Fig. 4), are anti-inflammatory and if N^1^-APA inhibits their activation it might promote inflammatory bowel disease like symptoms in the intestine). Along these same lines, our data potentially explain why the absence of detectable *speB* in certain species of bacteria was significantly associated with IBD in humans^24^ and that the loss of deoxyhypusine synthase in mice causes IBD-like pathologies in mice ^33^; both of these results could be explained by the accumulation of N^1^-APA in *speB* mutant bacteria which inhibits deoxyhypusine synthase function in animals. If true, our data suggest that existing inhibitors of the bacterial enzymes SpeA or SpeE could potentially be useful in the treatment of IBD because they would block N^1^-APA production. Future studies will likely be critical in determining if N^1^-APA contributes to IBD development in humans or if inhibiting N^1^-APA production in intestinal bacteria might alleviate IBD symptoms.

## Acknowledgements

We would like to acknowledge Craig Ellermeier for sharing *Bacillus* strains and plasmids. This research was supported in part by the Van Andel Institute Mass Spectrometry Core (RRID:SCR_024903; Grand Rapids, MI). We thank D. Chandler for helpful comments on the manuscript text, the *Caenorhabditis* Genetics Stock Center (funded by the NIH National Center for Research Resources) and National BioResource Project (NRBP) for *C. elegans* strains, and WormBase as a repository of *C. elegans* data. This work was supported by funds from the VAI MeNu research grant program and NIH grant DP2DK139569.

## Diversity and Inclusion Statement

All individuals involved in this study fulfilled the criteria for authorship required by Nature Portfolio journals and thus have been included as authors. Roles and responsibilities of all collaborators were agreed to in advance of research. All efforts were made to ensure no one involved in this study was hindered from participation based on background, ethnicity, or gender. This research also does not result in any known stigmatization, incrimination, discrimination or personal risk to anyone involved in this study.

## Author Contributions

K. Nauta, D. Gates, and N. Burton formulated the project. K. Nauta, D. Gates, M. Mechan, X. Wang performed the *C. elegans* experiments. K. Nguyen maintained *C. elegans* strains and provided support. C. Isaguirre, M. Gendjar, K. Nauta, and R. Sheldon developed and performed mass spectrometry experiments and data analysis. C. Krawczyk and M. Weiland formulated the cell culture experiments and M. Weiland performed the experiments and downstream procedures. K. Nauta, R. Sheldon, and N. Burton wrote the manuscript.

## DATA AVAILABILITY

All data reported in this paper are available in the Statistics Source Data.

## QUANTIFICATION AND STATISTICAL ANALYSIS

Ordinary one-way ANOVA p-value calculations followed by a Sidak’s multiple comparisons test were used to determine statistical significance for figures: 1D, 3A, 3C, 4B, and 4D. 2A and 2H statistics were calculated in Compound Discoverer 3.3.3.200. Ordinary one-way ANOVA p-value calculations followed by a Dunnett’s multiple comparisons test were performed to determine statistical significance for figures 3F. A Two-way ANOVA followed by a Tukey multiple comparisons test was performed to determine statistical significance for figure 3B. A Brown-Forsythe and Welch ANOVA with Dunnett’s T3 multiple comparisons correction was performed for figures 4F, 4G, and SFig. 7B. All calculations were performed in GraphPad Prism 10.1.2 (324) ^36^. No statistical method was used to predetermine sample size. The experiments were not randomized. The investigators were not blinded to allocation during experiments and outcome assessment.

## METHOD DETAILS

### *C. elegans* growth media and conditions

*C. elegans* strains were maintained at 15 °C or 20 °C on NGM agar plates (3 g/L NaCl, 17 g/L Agar, 2.5 g/L peptone (Bacto), 25 mM KPO_4_ (pH 6), 5 mg/L Cholesterol, 1 mM MgSO_4_, 1 mM CaCl_2_) seeded with *E. coli* OP50 (CGC, DA837). All *C. elegans* strains used in this study are listed in Table 3. *C. elegans* that were fed a diet of *B. subtilis* or *B. subtilis* mutants for experimentation were grown on modified NGM (3 g/L NaCl, 17 g/L Agar, 5.0 g/L peptone (Bacto), 25 mM KPO_4_ (pH 9.2), 5 mg/L Cholesterol, 1 mM MgSO_4_, 1 mM CaCl_2_).

**Table 3:**
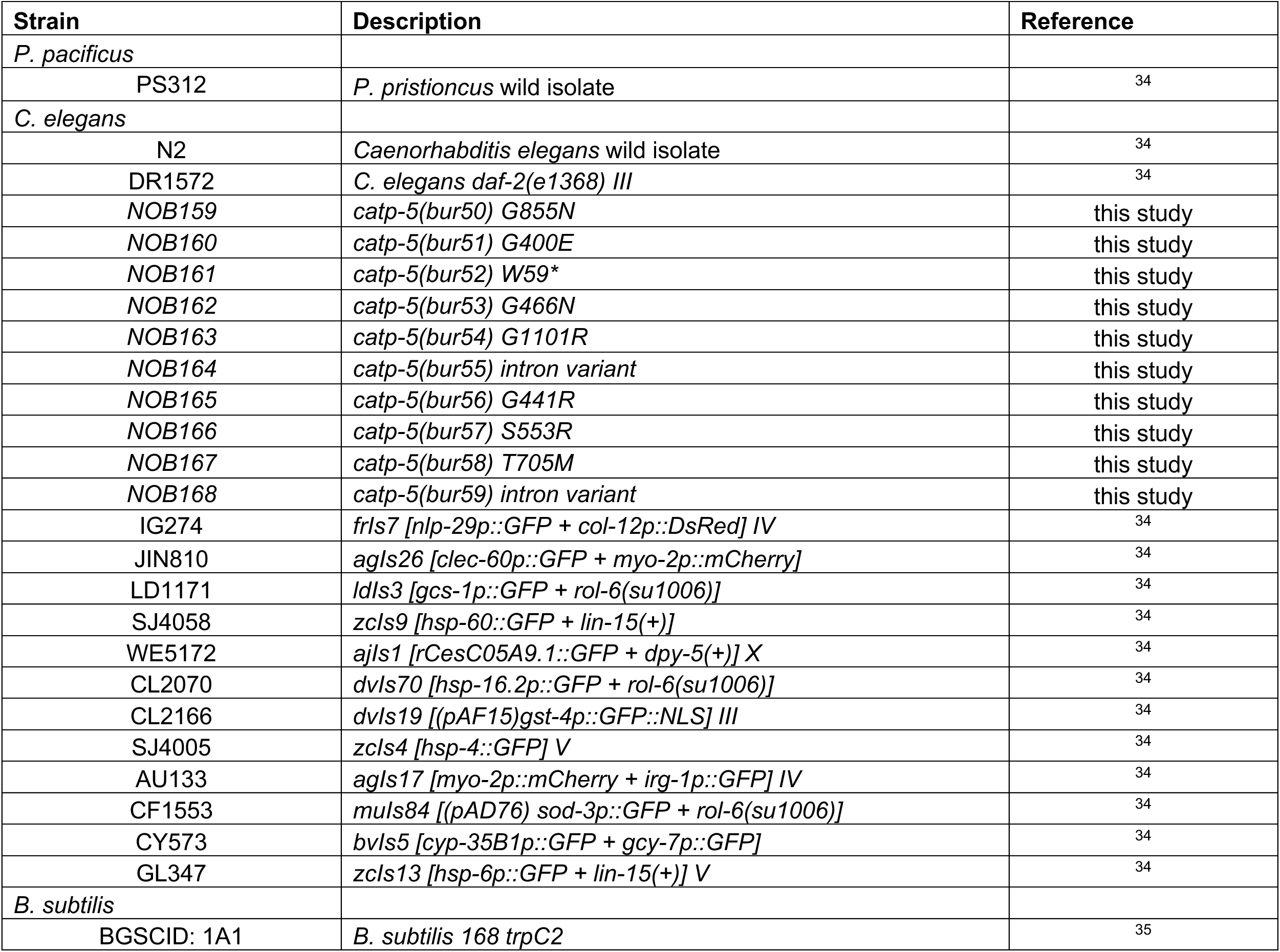
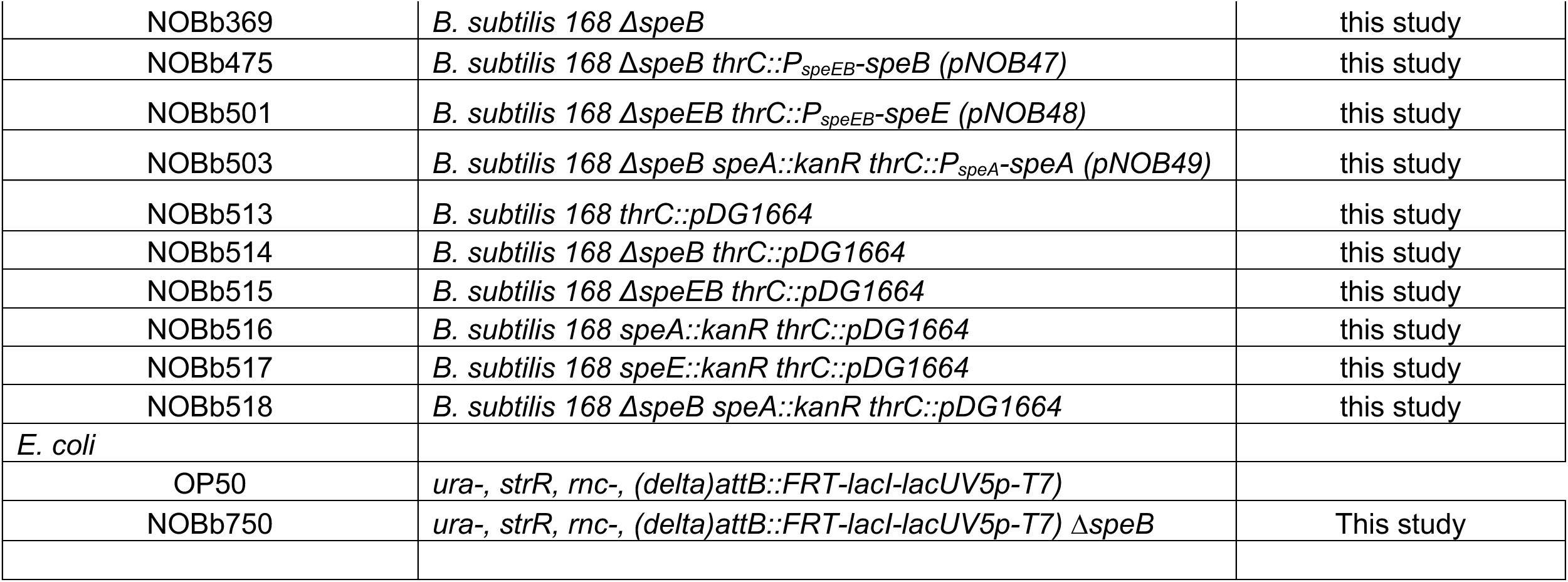
List of nematode and bacterial strains used in this study.

### Egg Prep Protocol

Adult *C. elegans* were washed from NGM agar plates with M9 buffer (3 g/L KH_2_PO_4_, 6 g/L Na_2_HPO_4_, 5 g/L NaCl, 1 mM MgSO_4_). *C. elegans* were lysed with Egg Preparation Solution (30% Sodium hypochlorite, 1 M NaOH) and monitored by dissection microscope for ∼^1-^3 minutes. The embryos were washed 2-3 times in M9 buffer.

### Bacteria growth media and conditions

All bacterial strains were grown on Lennox LB Agar (5 g/L NaCl, 10 g/L SELECT Peptone 140, 5 g/L SELECT Yeast Extract) (Invitrogen, 12780029) at 30 °C or 37 °C unless specified otherwise and are listed in Table 3. Liquid cultures were grown at 30 °C and shaken in <5 mL Lennox LB at 220 rpm in 18 mm glass test tubes. *Bacillus* strains containing pDG1664 and its derivatives inserted at *thrC* were grown in Lennox LB containing MLS (25 µg/mL erythromycin (Thermo Fisher Scientific, 50-213-296) and 5 µg/mL lincomycin (Thermo Fisher Scientific, 50-213-395)). Bacillus non-essential gene knock out library mutants were grown in kanamycin (7.5 µg/mL (Sigma, K1377)). Plasmid selection in *E. coli* was performed on Lennox LB containing kanamycin (25 µg/mL, Sigma Aldrich, K1377) or ampicillin (100 µg/mL, Sigma Aldrich, A0166).

### Bacteria strains and plasmid construction

All plasmids were constructed by isothermal assembly (ITA) and are listed in Table 4. DNA regions for plasmid assembly were identified using Biocyc ^37^ and amplified by touchdown PCR from the bacterial chromosomes using Q5 DNA polymerase (New England Biolabs, M0492) and oligonucleotides from Integrated DNA Technologies (Coralville, IA) listed in Table 5. All oligonucleotides are listed in Table 5. PCR fragments were purified using the Monarch DNA Gel Extraction Kit (New England Biolabs, T1020) and assemblies were performed using the NEBuilder HiFi DNA Assembly Master Mix (New England Biolabs, E2621). All regions amplified by PCR were confirmed by DNA sequencing at Genewiz from Azenta Life Sciences (South Plainfield, NJ). Plasmids were transformed according to the manufacturer’s instructions and maintained in *E. coli* DH5α competent cells (New England Biolabs, C2988J). Plasmids or purified chromosomal DNA were transformed into *B. subtilis* strains by natural competency as previously described ^25^.

**Table 4:** List of plasmids used in this study.

| Plasmid | Relevant features | Parent vector | Restriction enzymes to digest parent vector | PCR primers | PCR template | Reference |
| --- | --- | --- | --- | --- | --- | --- |
| pDG1664 | *thrC, amp, erm, spec* | N/A | N/A | N/A | N/A | (54) |
| pMAD | *amp, erm, ori-pE194ts* | N/A | N/A | N/A | N/A | (55) |
| pNOB41 | pMAD Δ*speEB* | pMAD | BglII, EcoRI | KN429-KN432 | *B. subtilis* 168 | this study |
| pNOB47 | *thrC::P<sub>speEB</sub>-speB amp erm spec* | pDG1664 | BamHI, EcoRI | KN475-KN478 | *B. subtilis* 168 | this study |
| pNOB48 | *thrC::P<sub>speEB</sub>-speE amp erm spec* | pDG1664 | BamHI, EcoRI | KN492, KN493 | *B. subtilis* 168 | this study |
| pNOB49 | *thrC::P<sub>speA</sub>-speA amp erm spec* | pDG1664 | BamHI, EcoRI | KN490, KN491 | *B. subtilis* 168 | this study |

**Table 5:**
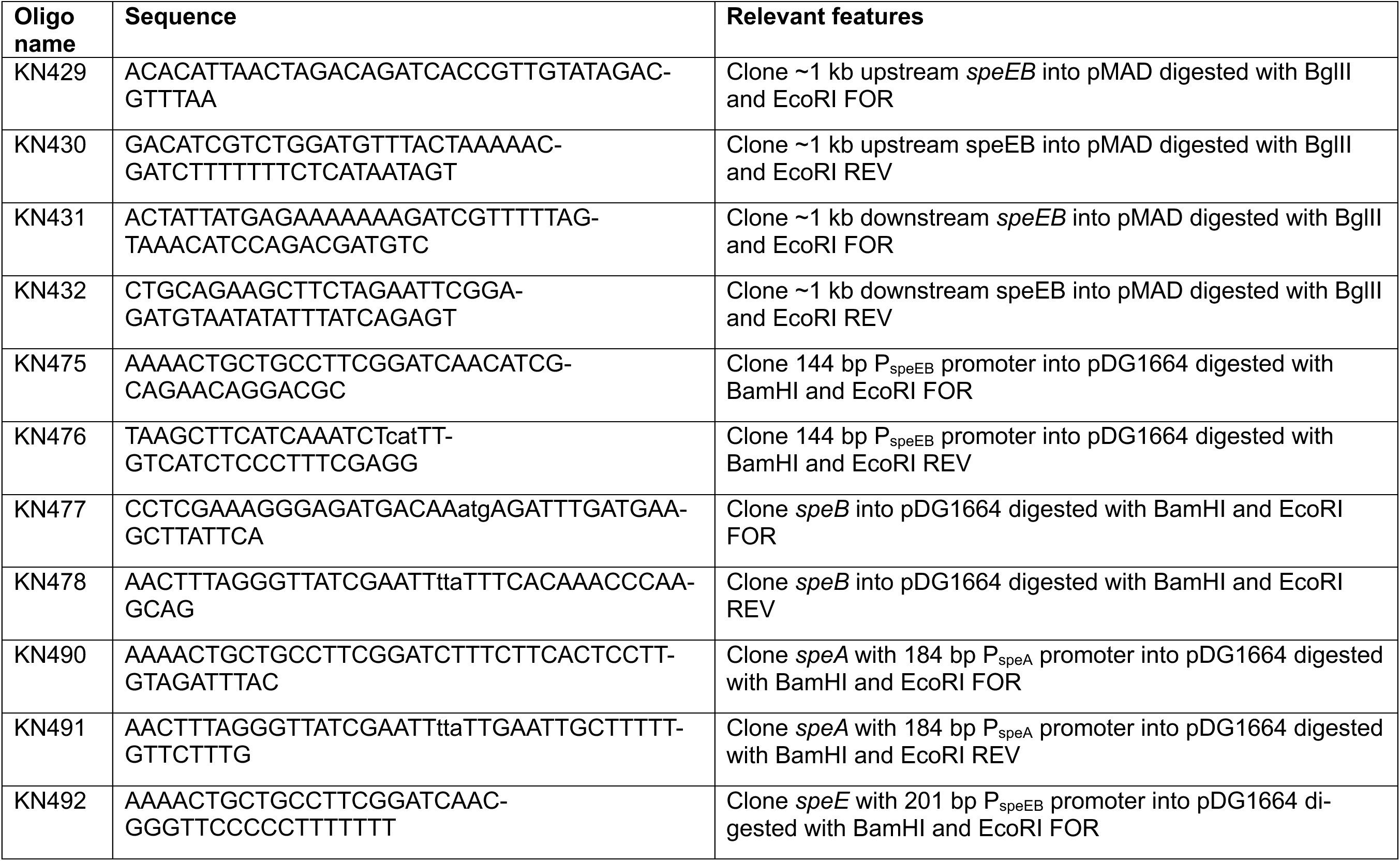
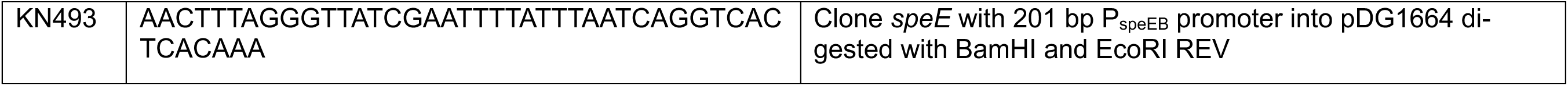
List of oligonucleotides used in this study.

All *Bacillus subtilis* strains reported in this study are isogenic derivatives of *B. subtilis* 168 (BGSCID: 1A1) and are reported in Table 3. The operon encoding *speEB* was knocked out by homologous recombination using the suicide vector, pMAD ^38^. Approximately 1 kb upstream and downstream of *speEB* were amplified and assembled via ITA into pMAD digested with BglII and EcoRI (New England Biolabs, R0144 and R3101, respectively), resulting in plasmid pNOBb41 (Table 4). The deletion was confirmed by PCR, sanger sequencing, and whole genome sequencing.

Strain NOBb518 was constructed by transforming purified chromosomal DNA (see methods section: *Bacillus* genomic DNA extraction) from strain BKK14630 (*speA::kanR*) ^25^ into NOBb369 (Δ*speB*). Chromosomal DNA was transformed by natural competency as previously described ^25^.

Plasmids for complementation studies (pNOB47, pNOB48, and pNOB49) were constructed by amplifying target genes from the *Bacillus subtilis* 168 chromosomes using the primers listed in Table 4. These amplicons were assembled in pDG1664 digested with EcoRI and HindIII (New England Biolabs, R3101 and R3104, respectively) by ITA and were expressed under the control of their respective native promoters. When necessary *kanR* was removed from the *Bacillus subtilis* chromosome using pDR244 ^25^ (Bacillus Genetic Stock Center, BGSCID ECE274).

*E. coli* OP50 Δ*speB* (NOBb750) was constructed by P1 transduction using *E.coli* K12 Δ*speB* lysate. The lysate was prepared using *E. coli* P1 bacteriophage (ATCC, 25404-B1). Briefly, *E. coli* K12 Δ*speB* donor strain was grown overnight at 37 °C in LB (30 μg/mL kanamycin). The culture was diluted 1:100 in LB supplemented with 0.2% glucose and 5 mM CaCl_2_, and incubated at 37 °C for 45 min, 220 rpm. 100 μL of *E. coli* bacteriophage P1 (ATCC 25404B1^TM^) was added to the donor bacteria culture and incubated at 37 °C for 3 hr. 200 µL of chloroform were added and incubated at 37 °C for 5 min. The mixture was centrifuged at 9,200xg for 5 min at 4 °C. Lysate was further purified using a 0.45 μm filter before collecting the P1 phage-containing supernatant. Recipient strain *E. coli* OP50 was grown overnight at 37 °C in 3 mL LB broth. *E. coli* OP50 cells were resuspended in a P1 salt solution (10 mM MgCl_2_, 5 mM CaCl_2_). Next, 10 μL of the P1 lysate was mixed with 100 μL of the recipient cells and incubated with shaking (220 rpm) at 37 °C for 30 min. Then, 1 mL of LB and 200 μL of 1 M sodium citrate was added and incubated at 37 °C for 1 hr, before being centrifuged at 13,000xg for 2 min. Transformants were selected on kanamycin (30 µg/mL) and sodium citrate (5 mM sodium citrate) LB ^39,40^.

### Screen of wild bacterial isolates for *C. elegans* development

Compost samples from the Grand Rapids, MI area were collected, diluted in M9 buffer, filtered, and plated on LB. Single colonies were grown overnight at 30 °C and 200 µL of culture was seeded on NGMB 12-well plates. The plates were dried in a laminar flow hood. Approximately 100-200 germ-free *C. elegans daf-2* (DR1572) embryos were added to each well. The plates were incubated for 5 days at 26 °C. The abundance of L4+ animals was qualified by eye using a dissection microscope. Strains of bacteria that promoted *C. elegans* development were retested on 90 mm NGM plates to confirm.

### Screen of *B. subtilis* 168 knockout library for loss of *C. elegans* development

The *B. subtilis* single gene deletion library--kanamycin was acquired from Addgene (Watertown, MA, 1000000115). The parent strain was acquired from the Bacillus Genetic Stock Center (Columbus, OH, BGSCID: 1A1). The library was replicated into 96-well plates for further use. Each 96-well plate was replicated and incubated overnight at 30 °C in LB. Each well was seeded on NGMB in a 12-well plate and dried in a laminar flow hood. 50-100 *C. elegans daf-2* (DR1572) embryos were added to each well. The plates were incubated for 5 days at 26 °C. Each *B. subtilis* mutant that did not promote animal development was retested on a larger scale using 90 mm plates.

### *Bacillus* genomic DNA extraction and whole genome sequencing *– B. subtilis*

Single bacteria colonies were grown overnight in 3 mL liquid culture. Bacteria culture was centrifuged at 17,000 x g for 1 min. Genomic DNA was isolated as previously described (58). Briefly, the pelleted cells were resuspended in Chromo Lysis Buffer (50 mM EDTA, 0.1M NaCl, pH 7.5). Lysozyme was added (20 µL of 2.5 mg/mL) and incubated at 37 °C for 10 minutes. Sarkosyl was added (1%) and the cells were incubated for 5 minutes at 37 °C. DNA of each isolate was extracted with phenol/chloroform/isoamyl alcohol 25:24:1 and ethanol precipitation. Pelleted were dried and suspended in sterile DI water. Total genomic DNA was quantified using a qubit fluorometer and visualized on a 1% agarose gel for degradation. Genomic DNA from each isolate was submitted for whole genome sequencing on an Illumina platform PE150 at SeqCenter (Pittsburg, PA). FastQC version 0.12.1 (GPL v3) was used to assess the quality control of the raw sequencing data ^41^. Reads were trimmed using Trimmomatic version 0.39 ^42^ to remove adapter sequences and low-quality reads (Q<30). Unicycler version v0.5.0 with default parameters was used to assemble the bacterial genome ^43^.

### EMS mutagenesis of WT *C. elegans*

L4 staged WT *C. elegans* were washed in M9 buffer (recipe: 3 g/L KH_2_PO_4_, 6 g/L Na_2_HPO_4_, 5 g/L NaCl, 1 mM MgSO_4_) 3 times and resuspended in 4 mL M9 buffer. 20 µL of Ethyl methanesulfonate (EMS) (Sigma Aldrich, M0880) were added and the animals were incubated for 4 hours with a rotary mixer, at 20 °C. The animals were washed three times and plated in 20 pools of ∼200 on *E. coli* OP50 seeded NGM. The animals were screened on *B. subtilis ΔspeB* for development beyond L4. Approximately one animal from each pool that developed beyond L4 was isolated and whole genome sequenced.

### *C. elegans* genomic DNA extraction and whole genome sequencing

WT *C. elegans* were incubated on NGM for 3 days at 20 °C. The animals were washed in M9 buffer. and suspended 1:1 in Worm Lysis Buffer (50 mM KCl, 10 mM Tris (pH 8.3), 2.5 mM MgCl2, 0.45% Tween 20, 0.01% Gelatin, 0.45% Nonidet P 40 Substitute (Sigma Aldrich, 74385). The animals were incubated at 56 °C for 1 hour with 10 µL of proteinase K (20 mg/mL). After lysis, genomic DNA was purified by phenol chloroform extraction and ethanol precipitation. DNA pellets were resuspended in sterile DI water.

Whole genome Illumina sequencing and variant calling was performed by SeqCenter (Pittsburg, PA). Briefly, Illumina sequencing libraries were prepared using the tagmentation-based and PCR-based Illumina DNA Prep kit and custom IDT 10bp unique dual indices (UDI) with a target insert size of 320 bp. No additional DNA fragmentation or size selection steps were performed. Illumina sequencing was performed on an Illumina NovaSeq 6000 sequencer in one or more multiplexed shared-flow-cell runs, producing 2×151bp paired-end reads. Demultiplexing, quality control and adapter trimming was performed with bcl-convert1 (v4.1.5). The fastq paired end reads were prepared for variant calling using Samtools and then indexed using bwa-mem index. Raw reads were then mapped to the indexed reference using bwamem and subsequently sorted and converted to BAM files using Samtools. GATK’s3 MarkDuplicates functionality was used to remove duplicate reads from the alignment file. GATK’s Haplotype caller was then used to call variants on the alignment file. BcfTools was used to filter out variants with a QD < 2, or MQ < 40 or MQRankSum < −12.5, or ReadPosRankSum < −8, or FS > 60.0, or SOR >3.

### Microscopy

*C. elegans* strains were washed with M9 buffer prior to imaging to remove excess bacteria. The animals were paralyzed with tetramisole (∼10 µg/mL). and incubated for ∼2 minutes. Images were taken with a Leica DM6 B upright microscope. Images were pseudo-colored using FIJI ^44^. Images were assembled in Adobe Illustrator 2024 28.4.1 64-bit ^45^.

### Metabolomics Extraction

Metabolites from LB broth were extracted via the addition of ice cold 2:2:1 (v/v) mixture of acetonitrile:methanol:water (#A456, #A955, #W6, respectively; Thermo Fisher Scientific) at a ratio of 30 µl sample per mL of extraction solvent. The samples were vortexed for 10 sec, sonicated for 5 minutes in a water bath sonicator, and incubated on wet ice for 60 min. After incubation, samples were centrifuged at 17,000g and 4 °C to pellet the insoluble fraction. The metabolite-containing supernatant was collected, dried in a vacuum evaporator, and stored at - 80°C until derivatization.

### Polyamine Derivatization

Carbamylated-derivatives were generated by derivatizing the dried metabolite fraction with isobutylchloroformate. Dried metabolite extracts were resuspended in 200 µL of water containing 1 µg/mL of 1,6 Diaminohexane as an internal standard. 5 µL of sodium bicarbonate (1M, pH 9.0) was added to each sample vial followed by the addition 20 µL of isobutyl chloroformate (177989, Sigma). Samples were vortexed for 10 seconds then incubated at 37 °C for 15 minutes. After cooling to room temperature, 1 mL of diethyl ether (309966, Sigma) was added, and samples were vortexed for 10 seconds. Samples were then centrifuged for 1 minute to separate the organic and aqueous layers. 800 µL of the carbamylated metabolite containing organic layer was transferred to an autosampler vial, and subsequently dried under a gentle stream of nitrogen gas. Dried samples were resuspended in 100 µL of 1:1 (v/v) acetonitrile:water, vortexed, and 90 µL transferred to a fresh autosampler vial for LC-MS analysis. For LCMS analysis of N^1^-APA, chemical standard (EN300-47248193, Enamine) was prepared at 2 µg/mL in water and derivatized as described above as either a neat standard or spiked into dried metabolite extracts.

### Untargeted LC-MS

LC-MS analysis of samples for Fig. 2A-F was performed on derivatized metabolite extracts analyzed by high resolution accurate mass spectrometry using an ID-X Orbitrap mass spectrometer (Thermo Fisher Scientific) coupled to a Thermo Vanquish Horizon liquid chromatography system. 2 μL of sample volume was injected on column. Chromatographic separations were accomplished with a Cortecs T3 (120Å, 1.6 μm, 2.1 mm × 150 mm) analytical column (#186008500, Waters, Eschborn, Germany) fitted with a pre-guard column (120Å, 1.6 μm, 2.1mm × 5 mm; #186008508, Waters, Eschborn, Germany) using an elution gradient with a binary solvent system. Solvent A consisted of LC/MS grade water (W6–4, Fisher), and Solvent B was 99% LC-MS grade acetonitrile (A955, Fisher). Both mobile phases contained 0.1% (v/v) formic acid (A11710X1, Fisher). The 15-min analytical gradient at a flow rate of 400 μL/min was: 0–0.5 min hold at 100% A, 0.5–2.0 min ramp from 0% B to 40% B, 2.0-10.0 min from 40% B to 99% B, then held at 99% B from 10.0-15.0 min. Following the analytical separation, the column was re-equilibrated for 3.5 min as follows: 0–0.5 min hold at 100% A at 600 μL/min, 0.5–0.6 min decrease from 600 μL/min to 400 μL/min, 0.6–3.5 min hold at 100% A and 400 μL/min. The column temperature was maintained at 50°C. The H-ESI source was operated at spray voltage of 2500 V(negative mode)/3500 V(positive mode), sheath gas: 70 a.u., aux gas: 25 a.u., sweep gas: 1 a.u., ion transfer tube: 300°C, vaporizer: 250 °C. Sample data were collected via data dependent MS2 (ddMS2) fragmentation using MS1 resolution at 60,000, MS2 resolution at 7,500, intensity threshold at 2.0 × 104, and dynamic exclusion after two triggers for 10 s. MS2 fragmentation was completed first with HCD using stepped collision energies at 15, 30, and 45% and was followed on the next scan by CID fragmentation in assisted collision energy mode at 15, 30, and 45% with an activation Q of 0.25. MS1 scans used custom AGC target of 25% in positive mode and a maximum injection time of 50 ms. MS2 scans used standard AGC target and a maximum injection time of 22ms. The total cycle time of the MS1 and ddMS2 scans was 0.6 s. Untargeted metabolomics data were analyzed in Compound Discoverer (v 3.3, Thermo). Figures were prepared in GraphPad Prism and FreeStyle™ 1.8 SP2 QF1 Version 1.8.65.0.

### Targeted LC-MS

LC-MS analysis of samples for Fig. 2H was performed on derivatized metabolite extracts analyzed using an Agilent 6470 LC-QQQ mass spectrometer coupled to a Agilent 1290 UHPLC system as previously reported^21^. Targeted metabolomics data were analyzed in Skyline 23.1.0.455 (Agilent). Statistics were performed using ols and anova_lm from statsmodels 0.14.4, posthoc_ttest from scikit_posthocs 0.7.0 with no corrections, and multipletests, method = fdr_bh from statsmodels 0.14.4 for global p-value corrections. Figures were prepared in GraphPad Prism 10.3.0 (507).

### Brood size count

*Pristionchus pacificus* PS312 grew to adulthood on NGM plates seeded with *E. coli* OP50 at 20 °C. Embryos were isolated using egg preparation solution and plated on NGMB plates seeded with the respective bacterial strains. Bacterial strains were grown overnight at 30 °C before seeding. Once the animals reached the L4 growth stage, 10 animals were singled onto NGMB seeded with the respective bacterial strains, prepared from overnight cultures grown at 30 °C. The plates were incubated at 20 °C for the duration of the experiment. Offspring were counted each day for approximately 5 days. Animals that died on day 1 or 2 were removed from the data set.

### Cell Culture

Bone marrow cells from femur were isolated from C57BL/6 female mice, RBC lysed, pooled (n=3-6 mice), counted, and resuspended at 1×10^7/ml, and plated at 1.5×10^6 per well. To generate Bone Marrow Macrophages (BMMΦ) cells were cultured in RPMI supplemented with 10% FBS (Hyclone), 2mM L-glutamine (Gibco), 100U/mL penicillin/streptomycin (Gibco), with 20ng/ml M-CSF (R&D; 416-ML-050) for 7 days. BMM_Φ_(IL4) were generated with 20ng/ml IL-4 (Peprotech 214-14-20ug) overnight from day 7; N^1-^guanyl-1 7-diaminoheptane (Stock 20mM) (GC7 1:100) 200µM and N^1^-APA (Stock 20mM) (APA 1:100) 200uM treatment began on day 7.

### BMM Flow

Following treatment on D7, BMM_Φ_ were harvested, washed and stained for flow cytometry. Samples were incubated with Fc block and eFluor 506 Fixable Viability Dye (ThermoFisher) in PBS, followed by an antibody cocktail prepared in wash buffer (PBS with 1% FBS, 1 mM EDTA, and 0.05% NaN_3_). For intracellular staining, cells were fixed with IC fixation buffer (eBioscience/ThermoFisher) for 30 minutes, permeabilized using Permeabilization Buffer (eBioscience/ThermoFisher) and incubated for at least one hour with antibodies targeting intracellular proteins. Samples were acquired on the Cytek Aurora spectral cytometer and data analyzed using FlowJo v10.

### BMM_Φ_ Western Blot for Hypusine and eIF5a

After 24 hours of treatment, BMMΦ were removed from culture by washing with warm 1xPBS followed by a wash with chilled 1xPBS and placed on ice for 5 minutes then gently scraped, resuspended and put into 15ml falcon tubes. Cells were pelleted at 4°C for 5min at 400g. Pellets were lysed with 1x CHAPS lysis buffer (+PI;+PMSF) freeze thawed twice at −80°C, sonicated by 10 seconds in an ice slurry followed by 1 minute rest (repeated 5 times). Supernatant was clarified by centrifuging for 15 minutes 4°C at 17,000xg. Lysed samples were stored at −80°C. Protein quantification was done with the Pierce 660nm Protein Assay Reagent (ThermoFisher; 22660); samples were diluted 1:2 in 1xCHAPS and read on a plate reader along with Pre-diluted Protein Assay Standards: Bovine Serum Albumin Set (ThermoFisher; 23208). For Hypusination and eIF5A blots,1 to 5µg of protein for each sample was removed, brought up to volume with 1xCHAPS and 6xLB. Samples were heated at 70°C for 10 minutes. For Western Blot 4-20% mini-protean gels were used and ran at 70V for 30 minutes and 100V until desired (30min). Proteins were transferred to PVDF membranes, blocked for 1hr 4°C in 5% milk. Membranes were incubated with primary antibodies for anti-hypusine (1:10000) (Millipore ABS1064-I-25ul) and anti-eIF-5a (1:10000) (BD Biosciences 611976) overnight at 4°C in 5% BSA+TBST. Secondary antibodies, 1hr RT 1:10000 in 5% milk+TBST.

---

**Supplementary Figure 1:**
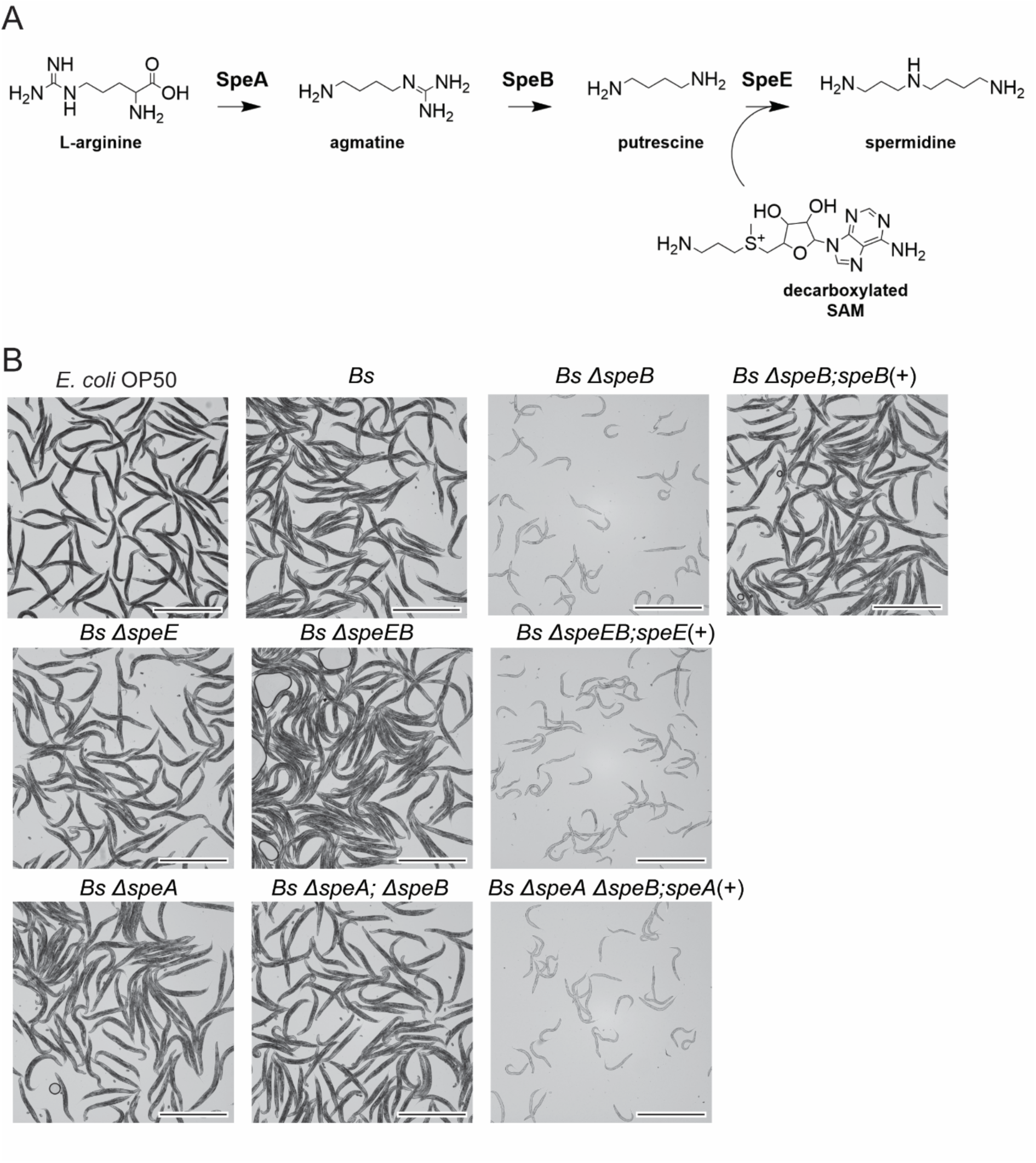
Loss of B. subtilis *speB* causes *C. elegans* developmental arrest. (A) Canonical polyamine pathway. (B) Representative images corresponding to Fig. 1D. Scale bars = 1 mm.

**Supplementary Figure 2:**
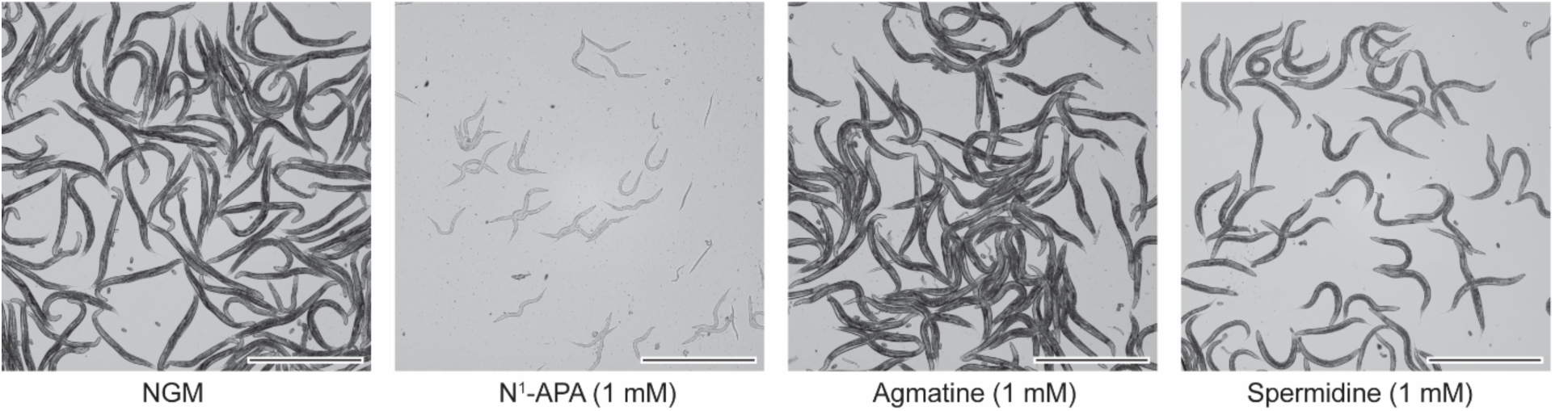
Exogenous spermidine and agmatine do not cause developmental arrest. Representative images corresponding to Fig. 3A of WT *C. elegans* grown on NGM seeded with *B. subtilis* WT (0) and supplemented with N^1^-APA (1 mM), agmatine (1 mM), or spermidine (1 mM). Scale bar = 1 mm.

**Supplementary Figure 3:**
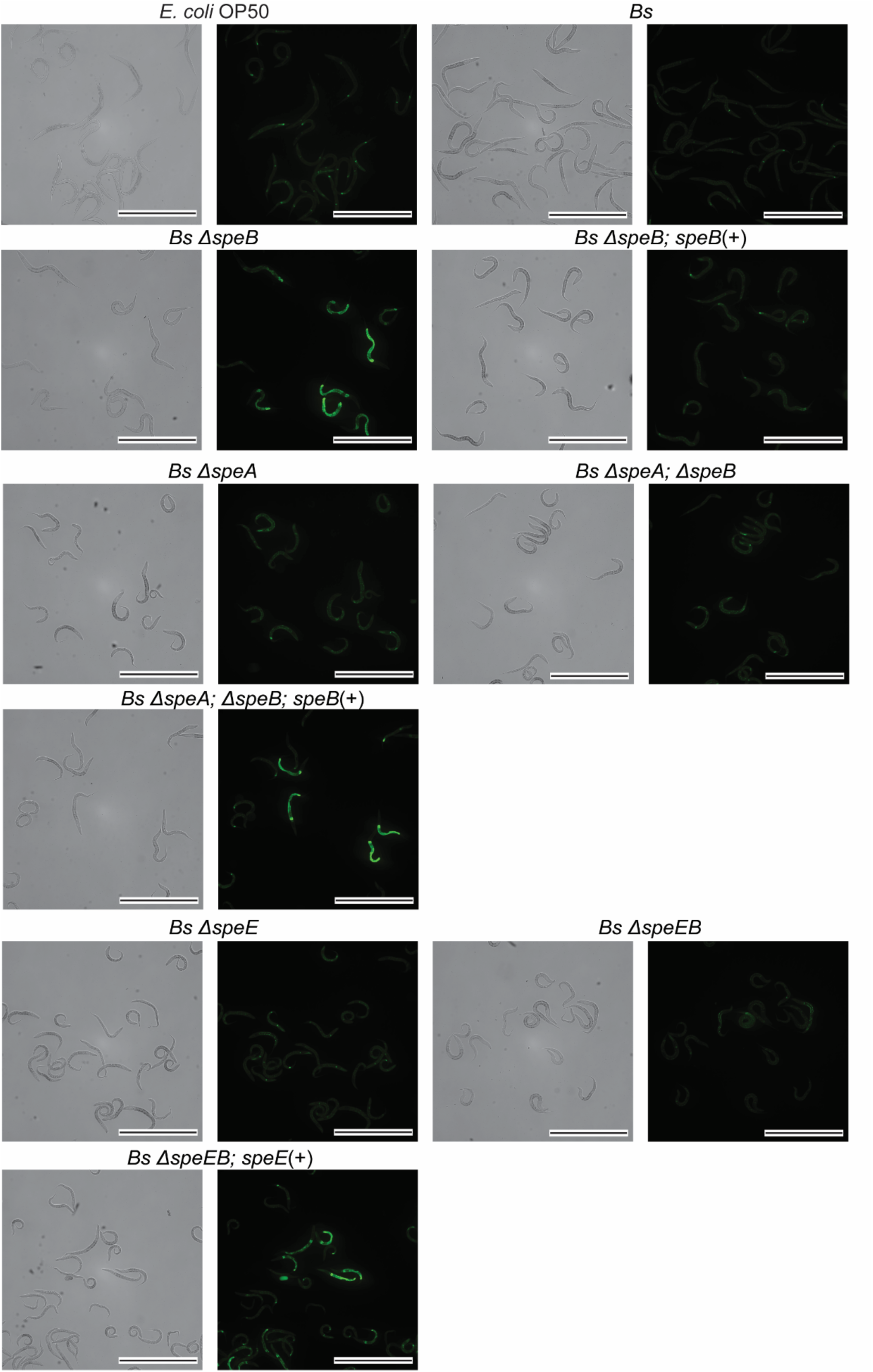
N^1^-APA-induced developmental arrest is mediated by mitochondrial stress. A) Brightfield (BF) and fluorescent (GFP) images of growth stage matched *hsp-6p::GFP*, on *E. coli* OP50, *B. subtilis* (NOBb513), *Bs* Δ*speB* (NOBb514), *Bs* Δ*speB; speB*(+) (NOBb475), (B) *Bs* Δ*speE* (NOBb517), *Bs* Δ*speEB* (NOBb515), *Bs* Δ*speEB; speE*(+) (NOBb501), *Bs* Δ*speA* (NOBb516), *Bs* Δ*speA; ΔspeB* (NOBb518), and *Bs* Δ*speA*; Δ*speB*; *speA*(+) (NOBb503). The *C. elegans* strains were incubated on the respective *B. subtilis* strains for ∼30 hours at 20 °C. Representative images were pseudo-colored using ImageJ v1.54f. Scale bars = 0.5 mm.

**Supplementary Figure 4:**
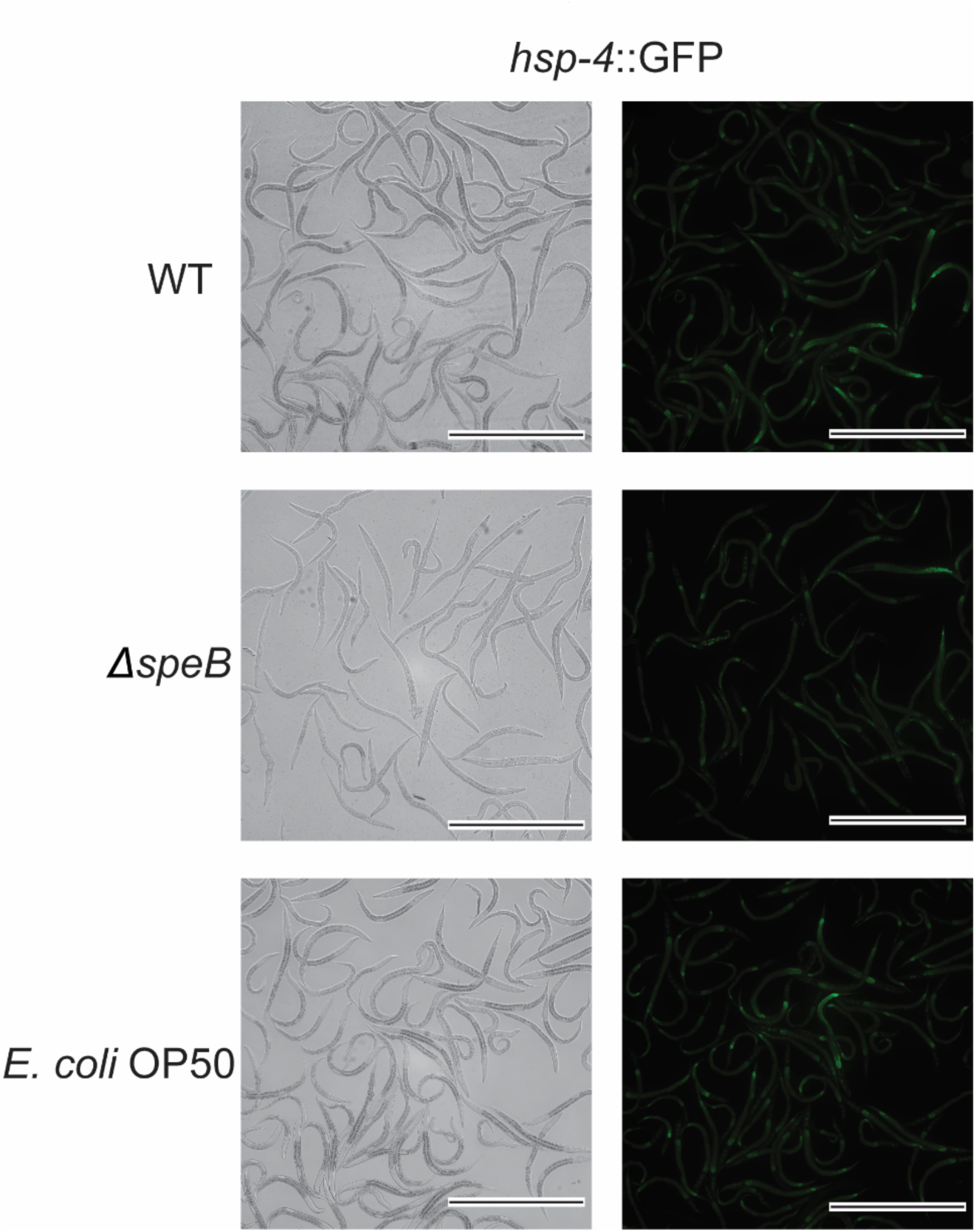
*hsp-4* is not activated by N^1^-APA. Brightfield (BF) and fluorescent (GFP) images of growth stage matched *hsp-4p::GFP*, on *E. coli* OP50, *B. subtilis* (WT), and *Bs* Δ*speB* (NOBb369). Representative images were pseudo-colored using ImageJ v1.54f. Scale bars = 0.5 mm.

**Supplementary Figure 5:**
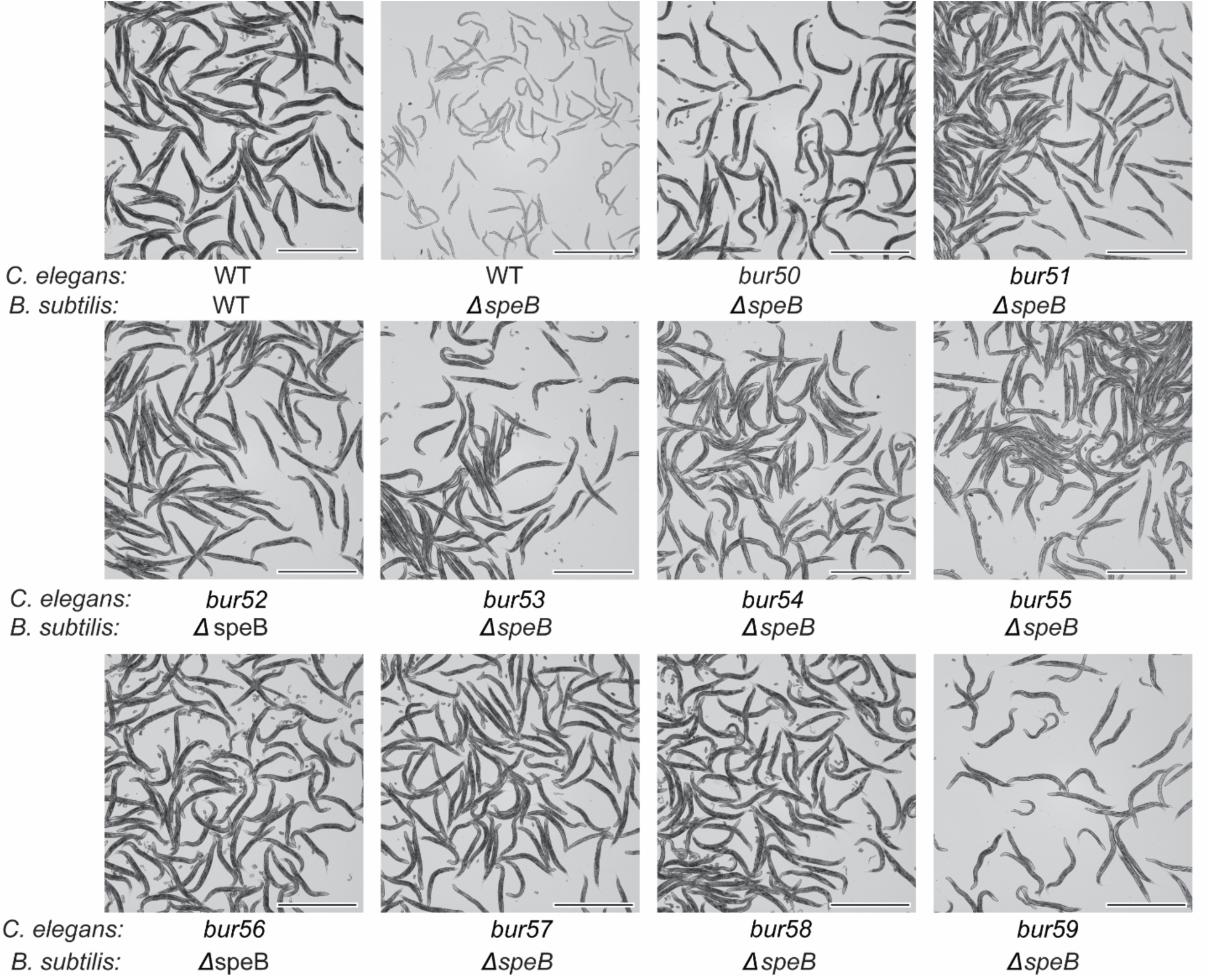
mutations in *catp-5* promote *C. elegans* development on a diet of *B. subtilis* Δ*speB*. Representative images of WT and *catp-5* (*bur50-59*) on a diet of *B. subtilis ΔspeB* (NOBb369) after incubation at 20 °C for 3 days. Scale bars = 1 mm.

**Supplemental Figure 6:**
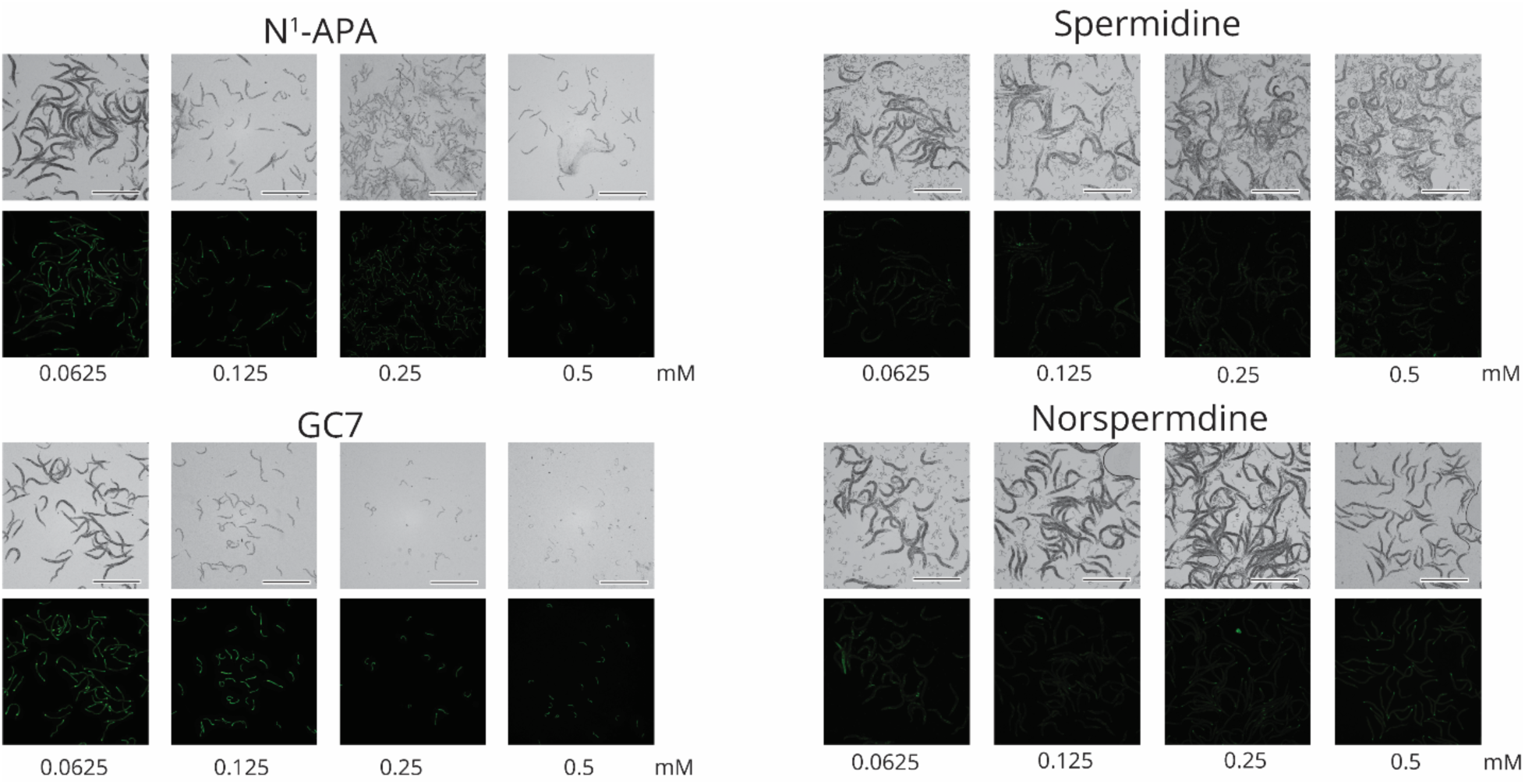
N^1^-APA and GC7 cause dose-dependent developmental arrest and expression of *hsp-6p::GFP*. Fluorescent (GFP) images of *C. elegans* encoding *hsp-6p::GFP* fed a diet of *B. subtilis* 168 on NGM agar plates supplemented with 0.0625-0.5 mM spermidine, norspermidine, N^1^-APA, and GC7. The *C. elegans hsp6p::GFP* reporter strain was exposed to the various polyamines for ∼30 hours at 20 °C on a diet of *B. subtilis* 168. Representative images were pseudo-colored using ImageJ v1.54f. Scale bars are 1 mm.

**Supplementary Figure 7:**
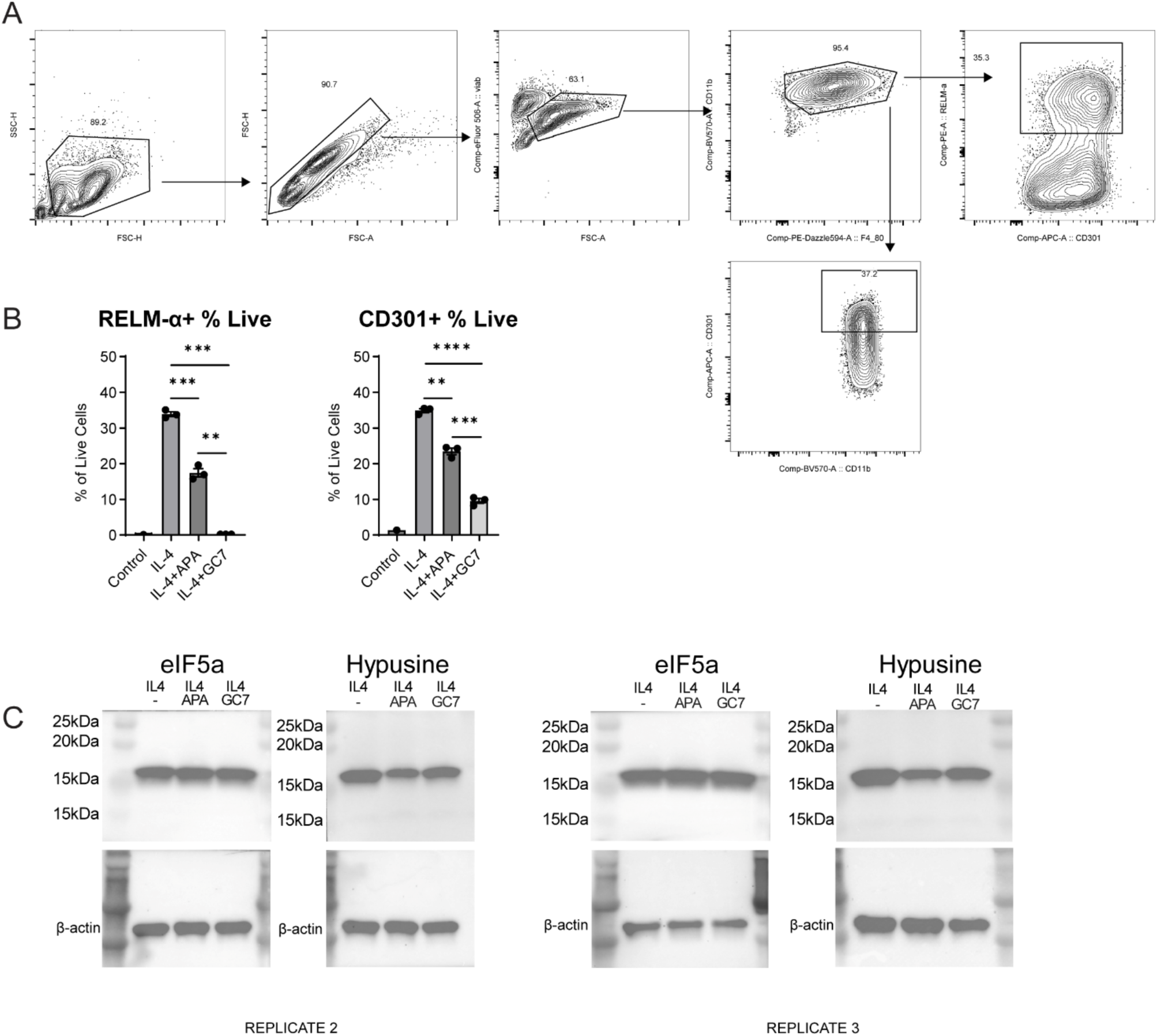
(A) Flow cytometry gating strategy corresponding to Fig. 4E. (B) Percentage of alive BMM_Φ_ after 24 hour treatment with 200 µM N^1^-APA (APA) or GC7 corresponding to Fig. 4F and 4G. Error bars = S.E.M., **** = p < 0.0001, *** = p < 0.009, ** = p <0.01. (C) Replicate western blots corresponding to Fig. 4H.

